# Autophagy disruption reduces mTORC1 activation leading to retinal ganglion cell neurodegeneration associated with glaucoma

**DOI:** 10.1101/2023.01.04.522687

**Authors:** Kang-Chieh Huang, Cátia Gomes, Yukihiro Shiga, Nicolas Belforte, Kirstin B. VanderWall, Sailee S. Lavekar, Clarisse M. Fligor, Jade Harkin, Adriana Di Polo, Jason S. Meyer

## Abstract

Autophagy dysfunction has been associated with several neurodegenerative diseases including glaucoma, characterized by the degeneration of retinal ganglion cells (RGCs). However, the mechanisms by which autophagy dysfunction promotes RGC damage remain unclear. Here, we hypothesized that perturbation of the autophagy pathway results in increased autophagic demand, thereby downregulating signaling through mammalian target of rapamycin complex 1 (mTORC1), a negative regulator of autophagy, contributing to the degeneration of RGCs. We identified an impairment of autophagic-lysosomal degradation and decreased mTORC1 signaling via activation of the stress sensor adenosine monophosphate-activated protein kinase (AMPK), along with subsequent neurodegeneration in RGCs differentiated from human pluripotent stem cells (hPSCs) with a glaucoma-associated variant of Optineurin (OPTN-E50K). Similarly, the microbead occlusion model of glaucoma resulting in ocular hypertension also exhibited autophagy disruption and mTORC1 downregulation. Pharmacological inhibition of mTORC1 in hPSC-derived RGCs recapitulated disease-related neurodegenerative phenotypes in otherwise healthy RGCs, while the mTOR-independent induction of autophagy reduced protein accumulation and restored neurite outgrowth in diseased OPTN-E50K RGCs. Taken together, these results highlight an important balance between autophagy and mTORC1 signaling essential for RGC homeostasis, while disruption to these pathways contributes to neurodegenerative features in glaucoma, providing a potential therapeutic target to prevent neurodegeneration.

## Introduction

Retinal ganglion cells (RGCs) are the sole projection neurons that connect the eye with the brain, and the degeneration of these cells in diseases such as glaucoma results in vision loss or blindness. Similar to other neurons throughout the central nervous system, RGCs are postmitotic and therefore highly dependent upon autophagy to remove damaged proteins or organelles to maintain proper cellular homeostasis. Autophagy deficits have been implicated in multiple neurodegenerative diseases including Alzheimer’s Disease, Parkinson’s Disease and Amyotrophic Lateral Sclerosis (ALS) (Barmada et al., 2014; Cuervo et al., 2004; Wong and Holzbaur, 2015; Yu et al., 2005). Similarly in glaucoma, a neurodegenerative disease characterized by the progressive degeneration of RGCs, the attenuation of autophagy had initially been observed in the trabecular meshwork and more recently described in RGCs (Hirt et al., 2018; Porter et al., 2015; Porter et al., 2014). In addition, a subpopulation of glaucoma patients possess mutations in the autophagy receptor Optineurin (OPTN) or the autophagy-associated protein TANK-binding kinase 1 (TBK1) resulting in glaucoma within a normal range of intraocular pressure, with the OPTN(E50K) mutation known to induce a severe degenerative phenotype (Aung et al., 2005; Rezaie et al., 2002; Ritch et al., 2014). Despite this, our knowledge of how autophagy impairment promotes neurodegeneration within RGCs remains limited.

The mechanistic target of rapamycin (mTOR) signaling pathway is a ubiquitous metabolic sensor, and signaling through mTOR complex 1 (mTORC1) is a well characterized primary modulator that negatively regulates autophagy processing (Boya et al., 2013). In the central nervous system, mTOR is responsible for neural stem cell proliferation, synaptic plasticity and neurite outgrowth, among other functions (Casadio et al., 1999; Lipton and Sahin, 2014; Tang et al., 2014; Tavazoie et al., 2005). In RGCs, activation of mTOR signaling either through the deletion of Phosphatase and tensin homolog (PTEN), a negative regulator of mTOR, or through the expression of Osteopontin, has been shown to promote axon regeneration after optic nerve injury (Duan et al., 2015; Park et al., 2008). Conversely, ocular hypertension induces hyperactivation of adenosine monophosphate-activated protein kinase (AMPK), resulting in dendrite retraction as well as synapse elimination via inhibition of mTORC1 in RGCs (Belforte et al., 2021). However, whether autophagy disruption affects mTOR signaling in RGCs is poorly understood, including the possibility that a greater demand for autophagy due to dysfunction of this pathway may negatively influence RGCs through the inhibition of mTOR signaling.

To address this shortcoming, in this study we have further advanced our hPSC-RGC neurodegeneration model with an underlying OPTN(E50K) mutation to study how autophagy disruption contributes to RGC neurodegeneration, and further validated these findings in a mouse ocular hypertension (OHT) glaucoma model with elevated intraocular pressure secondary to microbead occlusion. We identified a disruption of OPTN protein processing and a decrease in autophagic flux in hPSC-RGCs with the OPTN(E50K) mutation. Interestingly, OPTN(E50K) mutation-induced protein accumulation was observed in RGCs but not in related cortical neurons when derived from hPSCs, suggesting that dysfunction of OPTN function disrupts autophagy within OPTN(E50K) RGCs in a neuronal cell type-specific manner. More so, downregulation of mTORC1 signaling was observed in hPSC-RGCs with the OPTN(E50K) mutation, suggesting that autophagy disruption adversely affects other cellular pathways. To validate these results, an analysis of the mouse OHT model demonstrated an increase in the expression of autophagy proteins was associated with a decrease of mTOR signaling in RGCs, underscoring the concept that RGC neurodegeneration may be the result of imbalanced activity of mTOR-dependent autophagy. To test this idea, we found that deprivation of insulin, a positive regulator of mTOR signaling, exacerbated neurodegenerative phenotypes within hPSC-RGCs, while treatment with the mTOR-independent autophagy inducer trehalose rescued autophagy deficits and morphological phenotypes in hPSC-RGCs without affecting mTOR signaling. Thus, the results of these studies demonstrate that autophagy dysfunction promotes RGC neurodegeneration through inhibition of mTORC1 signaling, and that the modulation of these pathways can rescue neurogenerative phenotypes.

## Results

### The OPTN(E50K) mutation altered endogenous protein modification and resulted in accumulation of OPTN aggregates in hPSC-RGCs

We have previously established hPSC-derived RGCs carrying a BRN3b-tdTomato-Thy1.2 reporter as an in vitro neurodegenerative model through the use of CRISPR/Cas9 genome editing approaches to introduce the OPTN(E50K) glaucoma-associated mutation, causative for a severe neurodegenerative phenotype, into the otherwise unaffected H7 line of hPSCs (VanderWall et al., 2020). Similarly, we previously corrected the same mutation from a patient-specific induced pluripotent stem cell (iPSC) line with an OPTN(E50K) mutation (VanderWall *et al*., 2020). In the current study, we approached each set of analyses with these two isogenic pairs of cell lines including OPTN(E50K) and unaffected control (wild-type) hPSC-RGCs to minimize genetic variability among comparisons, and analyses were typically performed at stages of RGC maturation at which they had been previously demonstrated to exhibit morphological and functional disease-associated phenotypes (VanderWall *et al*., 2020).

To study how the endogenous E50K mutation attenuates RGC homeostasis contributing to neurodegeneration, we first characterized the expression of the autophagy receptor OPTN. While the level of *OPTN* mRNA was not significantly changed among wild-type and OPTN(E50K) RGCs, there was a 41.1 ± 5.2 % (mean ± S.E.M.) reduction in the level of OPTN protein in RGCs with the OPTN(E50K) mutation, along with a 28.1 ± 4.3 % increase in the autophagy receptor p62 (Figure 1A-C), consistent with previous findings that the E50K mutation decreased the overall abundance of OPTN protein in the mouse eye(Liu et al., 2021). To verify these results, we also reference our previously obtained RNA-seq data (VanderWall *et al*., 2020), which confirmed the trends observed in our qRT-PCR results. These results indicated that the E50K mutation likely altered OPTN protein during post translational modifications, and the increased level of p62 may be associated with autophagic accumulation. To rule out the possibility that the reduction of OPTN protein was due to a decreased ability of the antibody to recognize and bind to the E50K region, we performed an unbiased proteomics analysis of isogenic control and OPTN(E50K) RGCs two weeks after purification and identified 154 downregulated proteins as well as 178 upregulated proteins (Figure 1D). Among the downregulated proteins, OPTN identified with 4 peptides and 5 peptide-spectrum matches (PSM) was significantly decreased in OPTN(E50K) RGCs, corroborating our western blot results. Of interest, we identified an additional 7 autophagy-associated proteins whose expression was also altered, suggesting further disruption of the autophagy pathway due to the OPTN(E50K) mutation (Figure 1D).

**Figure 1.**
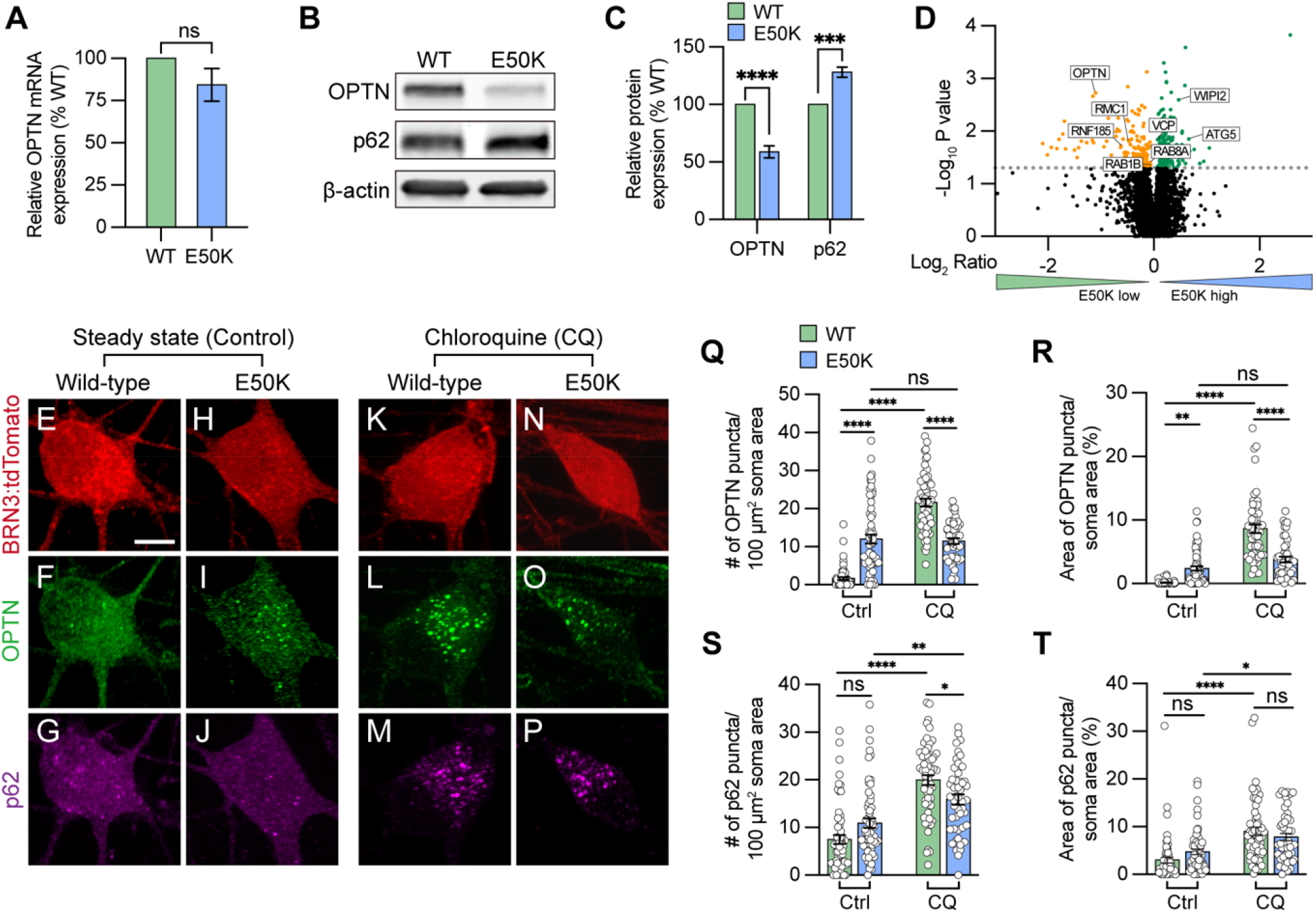
Modulation of OPTN levels in hPSC-derived OPTN(E50K) RGCs. (A) Real-time quantification of OPTN mRNA levels (n=3 for each WT and E50K; t-test, p=0.181). (B-C) Western blot and the relative protein expression of OPTN and p62 to β-actin in hPSC-RGCs (n=5 for each WT and E50K; t-test, OPTN ****p<0.0001, p62 ***p=0.0002). (D) Proteomics analysis demonstrated changes in the expression of autophagy-associated proteins in OPTN(E50K) hPSC-RGCs. (E-P) Immunostaining displayed the expression of OPTN and p62 puncta in BRN3:tdTomato hPSC-RGCs from WT and E50K under steady state (control) (E-J) and chloroquine (CQ) treatment (K-P). Scale bar: 10 μm. (Q-T) Quantification of OPTN puncta (Q and R) or p62 puncta (S and T) in hPSC-RGCs (n=3 biological replicates using Ctrl-WT n=60, Ctrl-E50K n=60, CQ-WT n=57 and CQ-E50K n=48 technical replicates; One-way ANOVA, Tukey post hoc test. ****p<0.0001, ***p<0.001, **p<0.01, *p<0.05, ns= not significant, p>0.05). Data are represented as mean values ± S.E.M. **Figure 1-source data 1. The alternation of OPTN and p62 protein in OPTN(E50K) mutation hPSC-RGCs**.

Under homeostatic conditions, the number of autophagosomes is balanced between autophagic biogenesis and degradation by the lysosome. Wild-type RGCs showed a predominantly cytosolic pattern with few puncta observed in RGC (Figure 1F) (Ying and Yue, 2012). However, we observed that OPTN(E50K) RGCs displayed a significant increase in OPTN puncta within RGC somas compared to wild-type RGCs, indicating an abnormal deposition of the OPTN protein in OPTN(E50K) RGCs, while no difference was observed in p62 puncta abundance between wild-type and OPTN(E50K) (Figure 1E-J and Q-T). To measure any preliminary changes in autophagic biogenesis without the loss of any autophagosomes due to lysosome-mediated degradation, we inhibited autophagosome-lysosome fusion by treatment with chloroquine for 16 hours prior to fixation (Klionsky et al., 2021). In wild-type RGCs, the number of OPTN and p62 puncta dramatically increased following chloroquine treatment, suggesting that autophagosome-lysosome mediated degradation was inhibited (Figure 1E-G, K-M, and Q-T). In contrast, the number of OPTN puncta in E50K RGCs did not change after treatment of chloroquine (Figure N, O, Q, and R), suggesting that autophagosome-lysosome mediated degradation was already impaired. Similar to wild-type RGCs, the number of p62 puncta was significantly increased in OPTN(E50K) RGCs after treatment with chloroquine, indicating that the E50K mutation did not affect p62 recruitment to the autophagosome (Figure 1J, P, S, and T). To rule out the possibility that the genomic background of the cell line caused this phenotype, including silent mutations introduced during CRISPR/Cas9 genome editing (VanderWall *et al*., 2020), we performed the same experiments and observed the same trends in patient-derived iPSC-derived RGCs harboring the OPTN(E50K) mutation in comparison with its CRISPR/Cas9-corrected isogenic control line (supplemental figure 1). Collectively, these findings suggest that the E50K mutation adversely affects RGCs due to the accumulation of OPTN protein and potentially contributes to RGC neurodegeneration.

### OPTN(E50K) hPSC-RGCs and mouse RGCs subjected to ocular hypertensive stress display autophagosome accumulation

During the process of autophagy, OPTN is necessary to recruit the microtubule-associated protein light chain 3 (LC3), for the engulfment of protein aggregates and/or damaged organelles, and for formation of the autophagosome (Evans and Holzbaur, 2020; Wong and Holzbaur, 2014). To study how the function of the OPTN protein changes due to the E50K mutation in RGCs, we initially used a GFP-fused-LC3 reporter to visualize the co-localization of LC3 and OPTN in RGCs. We found that 10.7 ± 1.2% of LC3 puncta co-localized with OPTN in wild-type RGCs, while only 5.1 ± 0.6% co-localized in RGCs with the OPTN(E50K) mutation (Figure 2A and B), suggesting that the E50K mutation attenuated OPTN recruitment of LC3. Inhibition of lysosome-mediated autophagosome degradation by chloroquine similarly demonstrated a decreased co-localization between OPTN and LC3 in OPTN(E50K) RGCs (supplemental figure 2). We further verified that the OPTN(E50K) mutation resulted in autophagy dysfunction by western blot. While there was no difference in the expression of cytosolic LC3-I in OPTN(E50K) RGCs (Figure 2C and D), significant increases were observed in the lipidated form of LC3-II, an indicator of autophagy (Kulkarni et al., 2020), as well as in autophagic flux as determined by the ratio of LC3-II/LC3-I (Figure 2D and E). Importantly, the level of the lysosomal protein LAMP1 also increased in OPTN(E50K) RGCs (Figure 2C and D), indicating that the OPTN(E50K) mutation not only induced autophagosome accumulation but also changed lysosomal degradation.

**Figure 2.**
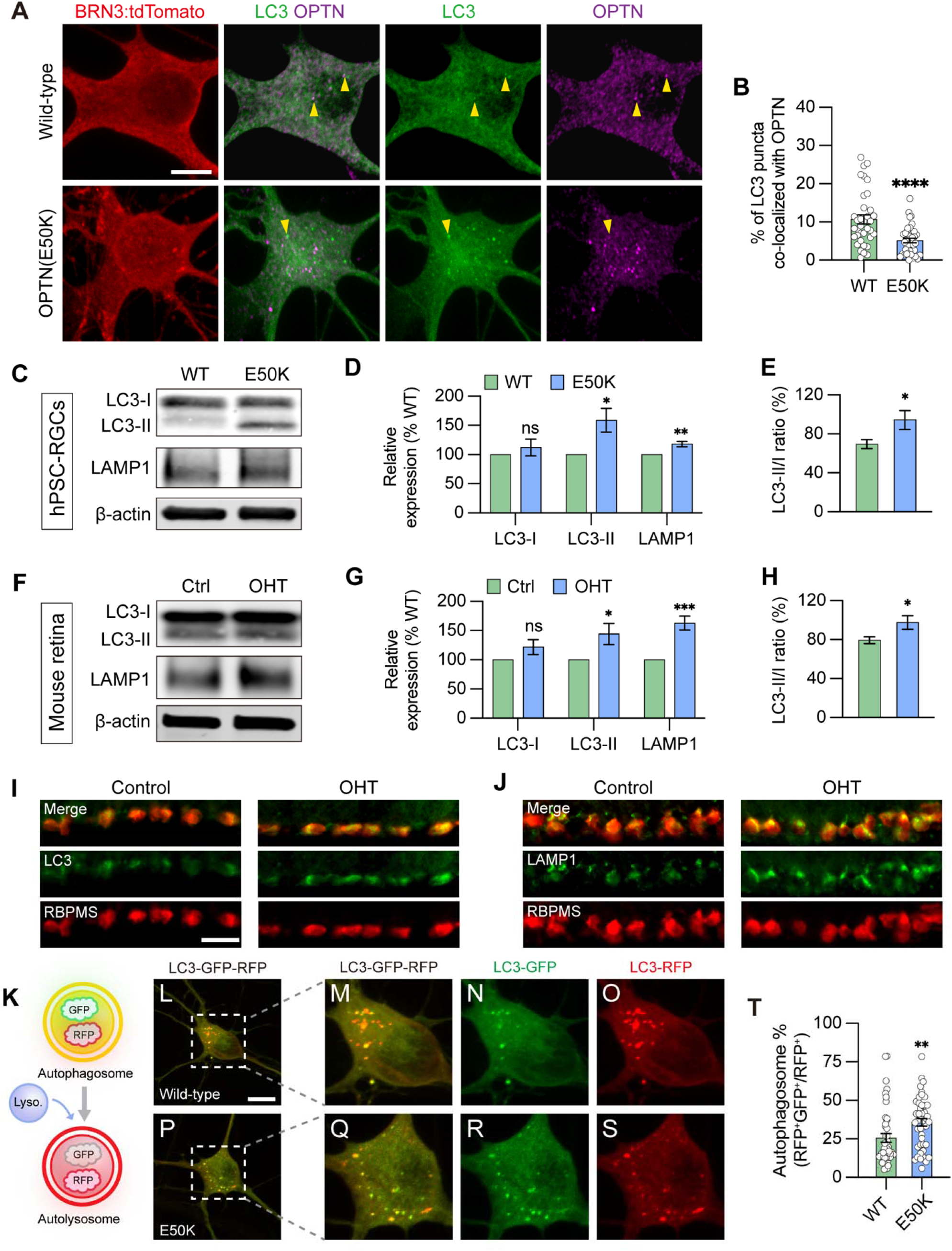
Disruption in autophagosome and lysosome degradation pathway in OPTN(E50K) hPSC-derived RGCs and ocular hypertensive mouse RGCs. (A-B) Analysis of protein colocalization between OPTN and LC3 in hPSC-RGCs (n=3 biological replicates using WT (n=36) and E50K (n=40); t-test, ****p<0.0001). Yellow arrows label the colocalization between OPTN and LC3 puncta. Scale bar: 10 μm. (C-E) Western blot and subsequent analysis of relative protein expression demonstrated increased LC3-II, LAMP1 and autophagic flux (LC3-II to LC3-I ratio) in OPTN(E50K) hPSC-RGCs (n=6 for each WT and E50K; t-test, LC3-I p=0.41, LC3-II *p=0.016, LAMP1 **p=0.0018, LC3-II/I *p=0.039). (F-H) Western blot verified changes in protein expression of LC3-I, LC3-II, LAMP1, and LC3-II/I ratio in control or glaucoma mouse retinas with ocular hypertension (OHT) (n=6 for each control and OHT; t-test, LC3-I p=0.122, LC3-II *p=0.034, LAMP1 ***p=0.0004, LC3-II/I *p=0.042). (I-J) Immunostaining displayed the elevation of LC3 and LAMP1 in RBPMS-expressing RGCs after ocular hypertension. Scale bar: 25 μm. (K) Schematic of LC3-RFP-GFP probe paradigm. (L-T) Using LC3-RFP-GFP probe showed the accumulation of autophagosome (RFP+GFP+) in OPTN(E50K) hPSC-RGCs (n=3 biological replicates using WT n=44 and E50K n=45 technical replicates; t-test, **p=0.0075). Scale bar: 10 μm. Data are all represented as mean values ± S.E.M. **Figure 2-source data 1. The alternation of LC3 and LAMP1 level in OPTN(E50K) mutation hPSC-RGCs**. **Figure 2-source data 2. The alternation of LC3 and LAMP1 level in mouse retina with ocular hypertension**.

Since multiple missense mutations in *OPTN* or *TBK1*, known as autophagy regulators, are known to result in normal tension glaucoma (Ritch *et al*., 2014; Swarup and Sayyad, 2018), we hypothesized that the maintenance of autophagy homeostasis plays a key role to maintain RGC survival and, conversely, impairment of autophagy may contribute to RGC degeneration in glaucoma. To further determine if RGC neurodegeneration is a consequence of autophagy dysfunction more broadly in glaucoma, we used a well-established magnetic microbead occlusion glaucoma model to induce ocular hypertension by injection of magnetic microbeads into the anterior chamber of the mouse eye. This procedure blocks aqueous humor outflow resulting in elevated IOP (supplemental figure 3) (Belforte *et al*., 2021; Ito et al., 2016), a major risk factor to develop glaucoma. Changes in autophagy markers were examined at 2 weeks after microbead injection, a time that precedes RGC loss thus avoiding the confounding effect of overt neurodegeneration (Belforte *et al*., 2021). In agreement with our findings in hPSC-RGCs, a significant increase in LC3-II, the LC3-II/I ratio, and LAMP1 were observed in the retina 2 weeks after ocular hypertension induction compared to sham-injected controls (Figure 2F-H). Immunostaining of retinal sections further showed the increased labeling of LC3 and LAMP1 more specifically in the ganglion cell layer, which co-localized with the RGC-specific marker RBPMS (Figure 2I and J). Collectively, our findings highlight that the disruption of autophagy is associated with RGC neurodegeneration not only in hPSC-RGCs with the OPTN(E50K) mutation, but more broadly in mouse RGCs in an ocular hypertension model, suggesting that autophagy disruption in RGCs may be a common mechanism across multiple glaucoma models.

### Autophagic-lysosomal degradation is impaired in OPTN(E50K) RGCs

A number of neurodegenerative diseases have been associated with a disruption to the autophagic-lysosomal pathway, including Alzheimer’s disease, Parkinson’s Disease, and ALS, leading to a deficit in protein degradation (Menzies et al., 2015; Nixon, 2013; Nixon and Yang, 2012). Since we observed a significant increase in the expression of autophagosome and lysosome proteins in OPTN(E50K) RGCs, as well as an increase in the LC3-II/LC3-I ratio, we further investigated whether these changes were specifically in response to disruption to the autophagic-lysosomal pathway by expressing RFP and GFP fused to LC3 in RGCs (Fig 2K), a widely used probe to determine autophagy flux (Kaizuka et al., 2016). This probe discriminated between the autophagosome and the acidic autolysosome due to differences in pH sensitivity, in which both the RFP and GFP signals were expressed in the autophagosome. While the RFP signal persisted, the GFP was extinguished under the acidic conditions within the autolysosome. As our previously established hPSC-RGC system included a BRN3-tdTomato reporter, whose red fluorescence could interfere with imaging using this probe, we used a similar approach to insert the OPTN(E50K) mutation in the H7 hPSC line lacking the BRN3-tdTomato reporter (VanderWall *et al*., 2020), and subsequently identified RGCs by staining with an antibody against BRN3 when imaging. In wild-type RGCs, 25.5 ± 2.8% of RFP puncta co-expressed GFP, indicating the remaining 74.5 ± 2.8% of autophagosomes had fused with the lysosome (Figure 2L-O, and T). However, OPTN(E50K) RGCs exhibited significantly more overlap between RFP and GFP at 35.8 ± 2.5%, representing an increase of 40.39% (Figure 2P-T), suggesting that the OPTN(E50K) mutation results in an impaired ability of the autophagosome to properly fuse with the lysosome for subsequent degradation.

### Selective degeneration of RGCs with the OPTN(E50K) mutation

To evaluate whether autophagy disruption selectively promotes RGC neurodegeneration rather than other cells that express the OPTN protein, we characterized relevant protein expression as well as neuronal morphologies in two projecting neurons including hPSC-RGCs and hPSC-cortical neurons (supplemental figure 4), respectively. We confirmed that differentiated wild-type and OPTN(E50K) cells expressed RGC identities based upon expression of BRN3 and MAP2, as well as cortical neuron identities based upon expression of CTIP2 and bIII-tubulin, respectively (Figure 3A and D). Concomitant with our previous findings (VanderWall *et al*., 2020), RGCs with the OPTN(E50K) mutation exhibited shorter neurites compared to wild-type, as analyzed by neurite complexity, soma size, number of primary neurites, and total neurite length (Figure 3B, C, G-I). In contrast, neurites from differentiated cortical neurons did not exhibit any significant differences between wild-type and OPTN(E50K) cell lines under the same measured parameters (Figure 3E-I), indicating that neurodegenerative features due to the OPTN(E50K) mutation were RGC specific. We further compared protein expression between RGCs and cortical neurons and identified that both neuronal types displayed a reduction in OPTN protein level due to the E50K mutation (Figure 3J and K). Interestingly, there was a higher cytosolic LC3-I expression in RGCs compared to cortical neurons (Figure 3J, lane 1 and 3), and only OPTN(E50K) RGCs exhibited a significant increase in the lipidated form of LC3-II (Figure 3J, L, and M). More so, unlike RGCs (Figure 1E-J), OPTN puncta did not accumulate in cortical neurons with the OPTN(E50K) mutation, comparable to wild-type cortical neurons during steady state conditions (Figure 3N-Q, V, and W). However, consistent with changes in RGCs (Figure 1K-P), the E50K mutation reduced total OPTN puncta in cortical neurons under chloroquine treatment in agreement with western blot results (Figure 3R-W), while the abundance of p62 puncta was not affected in cortical neurons with the OPTN(E50K) mutation (supplemental figure 5). Collectively, our findings suggest that the OPTN(E50K) mutation selectively renders RGCs more vulnerable to neurodegeneration because the higher demand for autophagy in RGCs is not met effectively.

**Figure 3.**
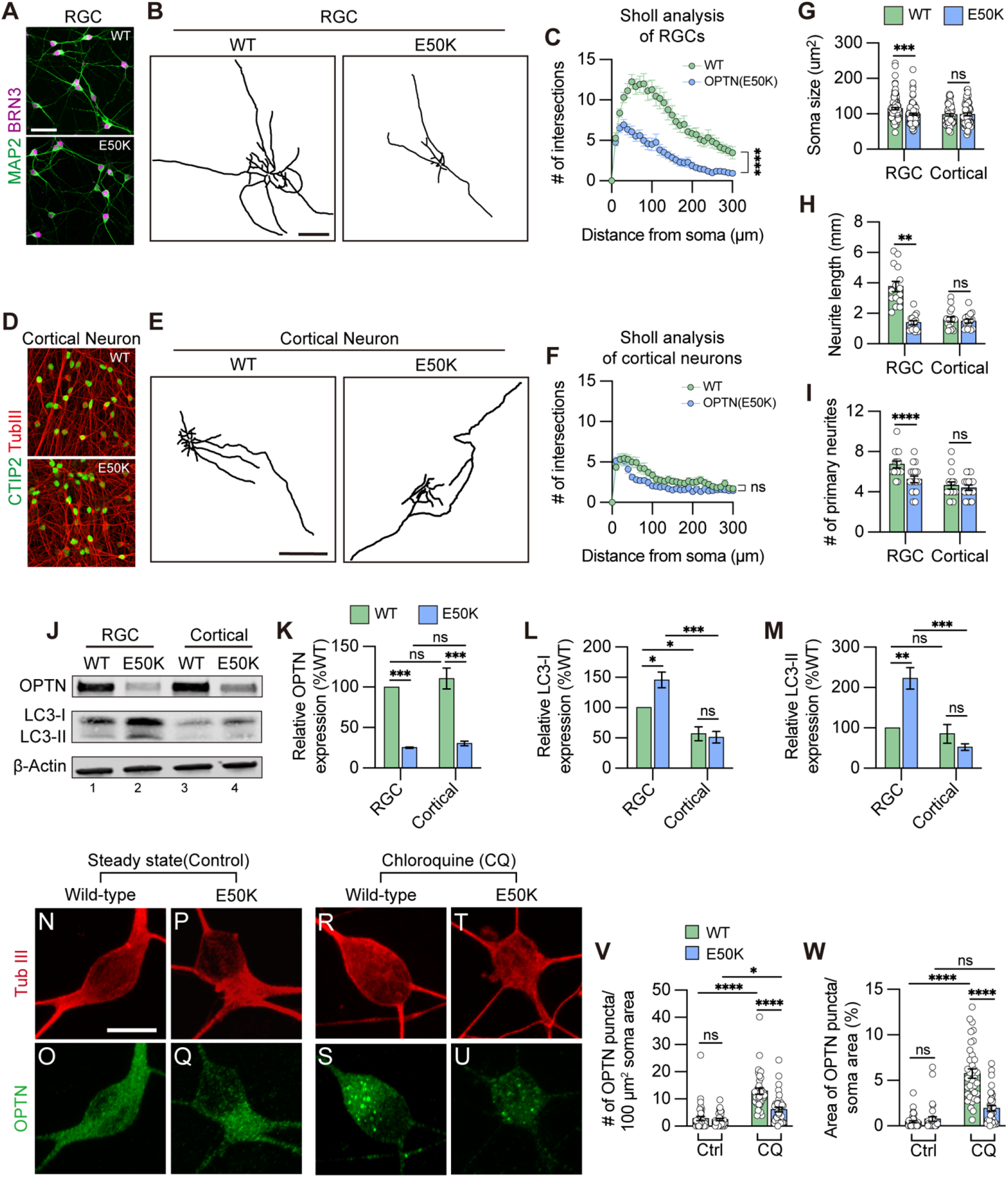
OPTN(E50K) mutation-induced phenotypes was tissue specific in RGCs. (A) Immunostaining to characterize wild-type and OPTN(E50K) RGCs. Scale bar: 25 μm. (B) Representative neurite tracing of wild-type and OPTN(E50K) RGCs after 4 weeks of purification. Scale bar: 150 μm. (C) Sholl analysis revealed the neurite complexity in wild-type and OPTN(E50K) RGCs (n=3 biological replicates using WT n=15 and E50K n=17 technical replicates; t-test, ****p<0.0001). (D) Immunostaining to characterize wild-type and OPTN(E50K) cortical neurons. Scale bar: 25 μm. (E) Representative neurite tracing of wild-type and OPTN(E50K) cortical neurons 4 weeks after purification. Scale bar: 150 μm. (F) Sholl analysis revealed similar neurite complexity in wild-type and OPTN(E50K) cortical neurons (n=3 biological replicates using WT n=16 and E50K n=17 technical replicates; t-test, ns=not significant, p=0.37). (G-I) Quantitative analysis of neurite parameters displayed neurite deficits in OPTN(E50K) RGCs based upon measurements of soma size (n=3 biological replicates using RGC-WT n=101, RGC-E50K n=112, cortical-WT n=68, and cortical-E50K n=66 technical replicates; t-test, RGC: *** p=0.0003; cortical: ns= not significant, p=0.811) (G), number of primary neurites (n=3 biological replicates using RGC-WT n=15, RGC-E50K n=17, cortical-WT n=16, and cortical-E50K n=17 technical replicates; t-test, RGC: ** p=0.004; cortical: ns= not significant, p=0.626) (H) and total neurite length (n=3 biological replicates using RGC-WT n=15, RGC-E50K n=17, cortical-WT n=16, and cortical-E50K n=17 technical replicates; t-test, RGC: **** p<0.0001; cortical: ns= not significant, p=0.575) (I), but not in OPTN(E50K) cortical neurons when compared with wild-type. (J-M) Western blot and quantified relative protein expression demonstrated that the OPTN(E50K) mutation altered LC3-II only in RGCs (n=3 for each WT and E50K from RGC or cortical neurons; One-way ANOVA, Tukey post hoc test. ***p<0.001, **p<0.01, *p<0.05, ns= not significant, p>0.05). (N-U) Immunostaining displayed the expression of OPTN puncta in hPSC-derived cortical neurons from WT and E50K under steady state (control) (N-Q) and chloroquine (CQ) treatment (R-U). Scale bar: 10 μm. (V-W) Quantification of OPTN puncta in hPSC-derived cortical neurons (n=3 biological replicates using Ctrl-WT n=39, Ctrl-E50K n=34, CQ-WT n=37 and CQ-E50K n=37 technical replicates; One-way ANOVA, Tukey post hoc test. ****p<0.0001, ***p<0.001, *p<0.05, ns= not significant, p>0.05). Data are all represented as mean values ± S.E.M. **Figure 3-source data 1. Western blot analysis in OPTN and LC3 proteins in hPSC-RGCs and cortical neurons**.

### Autophagy dysfunction results in downregulation of mTORC1 signaling

The mammalian target of rapamycin (mTOR) signaling pathway is a key metabolic regulator and sensor of stress. Activation of mTOR is known to promote dendritic morphological complexity as well as induce axonal regeneration in RGCs (Proietti Onori et al., 2021), while also functioning as a negative regulator of autophagy through the mTORC1 complex (Agostinone et al., 2018; Duan *et al*., 2015; Ganley et al., 2009; Hosokawa et al., 2009). Under cellular stress in neurons, autophagy disruption can activate adenosine monophosphate-activated protein kinase (AMPK) to induce the repression of mTORC1, resulting in neurodegeneration (Jung et al., 2019; Yang et al., 2020). To determine whether autophagy disruption promotes RGC degeneration via inhibition of mTORC1 signaling, we used pS6^Ser240/244^ as a readout of mTORC1 activity. mTORC1 induces the p70 ribosomal S6 kinase activation and subsequently phosphorylates the ribosomal protein S6 at Ser240/244 residues (pS6^Ser240/244^) (Jefferies et al., 1997). Indeed, RGCs from sham-injected control mice exhibited robust mTORC1 activity, as visualized by pS6^Ser240/244^ co-localization with the RGC marker RBPMS, while decreased mTORC1 activity was observed in RGCs subjected to ocular hypertensive stress (Figure 4A-E). This observation is in agreement with previous findings that mTORC1 signaling is partially inactivated in glaucomatous RGCs through AMPK phosphorylation resulting in dendritic retraction (Belforte *et al*., 2021). To evaluate whether the changes in mTORC1 could be observed in OPTN(E50K) hPSC-RGCs, we first characterized mTORC1 activity in hPSC-derived retinal cells. We analyzed cells isolated from retinal organoids after 50 and 80 days of differentiation, respectively, which allowed for the analysis of the majority of neuroretinal cell types including RGCs (BRN3B-tdTomato), retinal progenitors (CHX10), and photoreceptors (OTX2) (supplemental figure 6A-C). Immunohistochemistry detection of pS6^Ser240/244^ in hPSC-derived retinal cells revealed robust expression within BRN3B:tdTomato RGCs, but not CHX10-expressing retinal progenitor cells nor OTX2-positive photoreceptors, indicating a strong role for mTOR signaling specifically within hPSC-RGCs (supplemental figure 6D and E). Subsequently, we analyzed RGCs that were isolated from retinal organoids and allowed to mature for an additional 4 weeks, a timepoint at which we have previously demonstrated to result in some neurodegenerative phenotypes in OPTN(E50K) RGCs (Figure 3B), including neurite retraction and hyperexcitability (VanderWall *et al*., 2020). These studies revealed that OPTN(E50K) RGCs exhibited a decrease in the expression of the mTORC1 effector pS6^Ser240/244^, while no difference was observed in the expression of the mTORC2 effector pAKT^Ser473^ (Figure 4F and G). More so, OPTN(E50K) RGCs also increased the expression of the phosphorylated form of AMPK (pAMPK^Thr172^), which is activated under stress and functions to inactivate mTORC1 (Figure 4H). Immunohistochemistry also revealed a profound decrease in the expression of pS6^Ser240/244^ intensity in the somatic area of OPTN(E50K) RGCs (Figure 4I-O). Taken together, our results suggest that chronic autophagy deficits in glaucomatous RGCs activate AMPK signaling to suppress mTORC1, leading to neurodegeneration.

**Figure 4.**
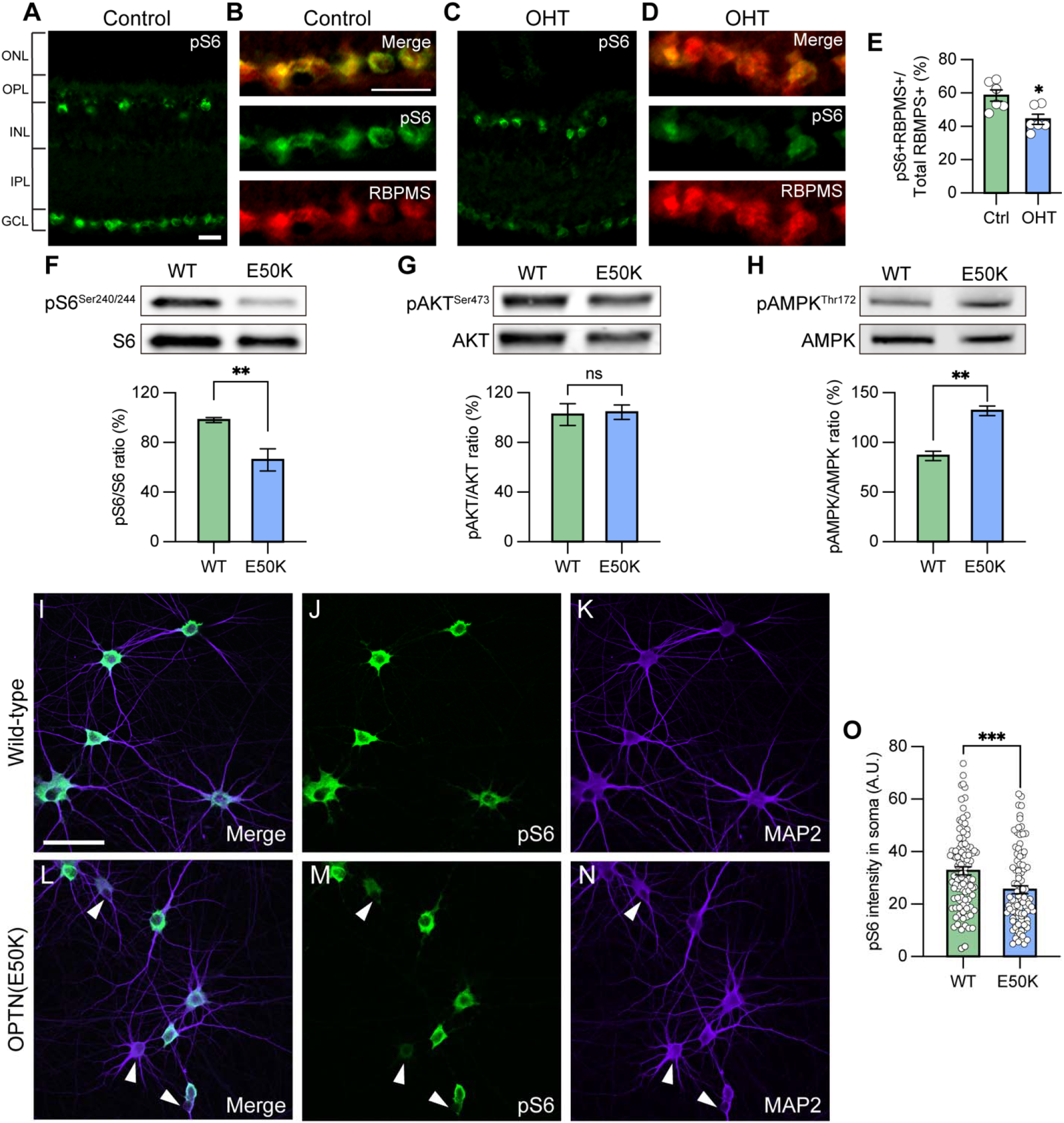
Attenuation of mTORC1 signaling in OPTN(E50K) hPSC-derived RGCs and ocular hypertension mouse RGCs. (A-B) Immunostaining labeled the level of the mTORC1 effector pS6^Ser240/244^ and exhibited robust activity in the ganglion cell layer (GCL), co-labeled with RBPMS in (B), and inner nuclear layer (INL) in control mouse retina. Scale bar: 25 μm. (C-E) Under ocular hypertension, immunostaining and associated quantification demonstrated that the level of pS6^Ser240/244^decreased in mouse RGCs (n=3 mice/group, 2 images/mice; t-test, *p=0.011). (F-H) Western blot and the relative protein expression of pS6^Ser240/244^, pAKT^Ser473^ and pAMPK^Thr172^ to its total protein, respectively, in hPSC-RGCs (n=6 for each WT and E50K; t-test, pS6^Ser240/244^**p=0.0057; pAKT^Ser473^ ns= not significant, p=0.854; pAMPK^Thr172^ **p=0.0027). (I-O) Immunostaining and quantification of pS6^Ser240/244^ intensity revealed that a subset of RGCs with the OPTN(E50K) mutation reduced the expression of pS6^Ser240/244^ (n=3 biological replicates using WT n=96 and E50K n=96 technical replicates; t-test, ***p=0.0009). Scale bar: 25 μm. Data are all represented as mean values ± S.E.M. **Figure 4-source data 1. The level of phosphorylated form S6 (Ser240/244) protein in hPSC-RGCs**. **Figure 4-source data 2. The level of phosphorylated form AKT (Ser473) protein in hPSC-RGCs**. **Figure 4-source data 3. The level of phosphorylated form AMPK (Thr172) protein in hPSC-RGCs**.

### Inhibition of mTOR signaling results in neurodegenerative phenotypes in otherwise healthy hPSC-RGCs

We hypothesized that autophagy disruption due to the OPTN(E50K) mutation caused mTOR suppression, leading to neurodegenerative phenotypes in RGCs. To further evaluate whether decreased mTOR activity correlates with impaired RGC neurite outgrowth and autophagy modulation, we treated wild-type hPSC-RGCs with the dual mTORC1/2 inhibitor KU0063794 for one week (Figure 5A). RGCs exhibited a reduction of pS6^Ser240/244^ intensity following treatment with KU0063794 in a dose-dependent manner (Figure 5B and C). When treated with KU0063794 (1 μM), hPSC-RGCs revealed significant decreases in the expression of both mTORC1 and mTORC2 effectors pS6^Ser240/244^ and pAKT^Ser473^, respectively, while the level of pAMPK^Thr172^ was increased (Figure 5D and E). Furthermore, pharmacological inhibition of mTOR resulted in increased LC3-I and LC3-II, an indication of autophagy activation (Figure 5F and G). However, the level of autophagic flux (LC3-II/LC3-I) and lysosome protein LAMP1 did not change under mTOR inhibition when compared to vehicle control, suggesting that wild-type hPSC-RGCs can effectively balance cellular homeostasis between autophagy and acute mTOR inhibition. Analysis of RGC neurites demonstrated a decrease in neurite complexity that correlated with an increase in the concentration of KU0063794 (Figure 5H-L). As morphological features of RGCs were modulated by KU0063794 in a dose-dependent manner, these results suggest that the pathological hallmarks of neurite retraction observed in OPTN(E50K) hPSC-RGCs may be the result of decreased mTOR signaling.

**Figure 5.**
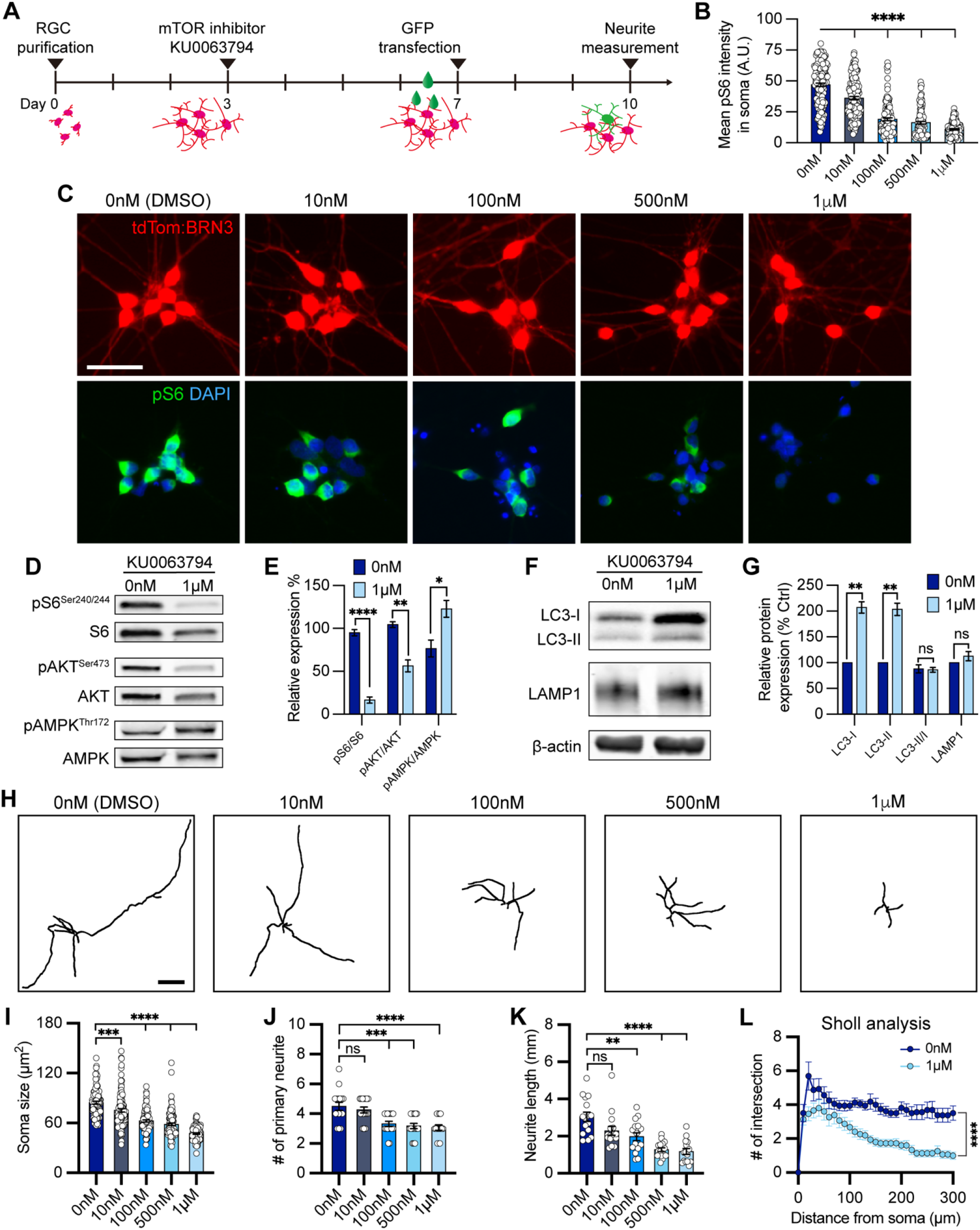
mTOR inhibition contributed to neurite shortening in hPSC-derived RGCs. (A) Schematic timeline of mTOR inhibitor treatment and methods for neurite analyses. (B-C) Quantification and immunostaining revealed that pS6^Ser240/244^ intensity was dose-dependent in response to mTOR inhibition in hPSC-RGCs (n=3 biological replicates using vehicle (DMSO) n=162, 10 nM n=143, 100 nM n=128, 500 nM n=121, and 1 μM n=133 technical replicates; One-way ANOVA, Tukey post hoc test. ****p<0.0001 for each KU0063794 treatment group compared to vehicle treatment). Scale bar: 25 μm. (D-G) Western blot and quantification of the relative protein expression displayed a decrease of mTOR signaling and induction of autophagy when mTOR was inhibited by KU0063794 treatment in hPSC-RGCs (n=3 for each vehicle (DMSO) and 1 μM of KU0063794 treatment, t-test, pS6^Ser240/244^ ****p<0.0001, pAKT^Ser473^ **p=0.0033, pAMPK^Thr172^ **p=0.0292, LC3-I ***p=0.0006, LC3-II **p=0.001, LC3-II/I p=0.838, LAMP1 p=0.23). (H) Representative neurite tracings of vehicle and KU0063794 treatment in hPSC-RGCs. Scale bar: 200 μm. (I-L) Quantitative analysis of neurite parameters displayed neurite deficits and decreased neurite complexity following mTOR inhibition in hPSC-RGCs as measured by their soma size (n=3 biological replicates using vehicle n=107, 10 nM n=109, 100 nM n=97, 500 nM n=107, and 1 μM n=93 technical replicates; One-way ANOVA, Tukey post hoc test. ****p<0.0001, ***p<0.001) (I), number of primary neurites (n=3 biological replicates using vehicle n=16, 10 nM n=16, 100 nM n=16, 500 nM n=17, and 1 μM n=15 technical replicates; One-way ANOVA, Tukey post hoc test. ****p<0.0001, ***p<0.001, ns= not significant, p>0.05) (J), total neurite length (n=3 biological replicates using vehicle n=16, 10 nM n=16, 100 nM n=16, 500 nM n=17, and 1 μM n=15 technical replicates; One-way ANOVA, Tukey post hoc test. ****p<0.0001, **p<0.01, ns= not significant, p>0.05) (K), and Sholl analysis (n=3 biological replicates using vehicle n=16, and 1 μM n=15 technical replicates; t-test, ****p<0.0001) (L). Data are all represented as mean values ± S.E.M. **Figure 5-source data 1. The level of phosphorylated form S6 (Ser240/244), AKT (Ser473), and AMPK (Thr172) proteins in hPSC-RGCs under mTOR inhibition**. **Figure 5-source data 2. The alternation of LC3 and LAMP1 level in hPSC-RGCs under mTOR inhibition**.

### Insulin deprivation exacerbates disease pathogenesis in OPTN(E50K) RGCs

Insulin is a canonical mTOR activator and previous studies have demonstrated the ability of insulin to rescue neurodegenerative phenotypes in dendritic arbors in rodent RGCs subjected to optic nerve crush (Agostinone *et al*., 2018). Previously, we have demonstrated that OPTN(E50K) RGCs exhibit morphological and functional deficits as soon as 4 weeks after the purification and maturation of RGCs from retinal organoids (Figure 3B) (VanderWall *et al*., 2020). However, as insulin is a common component of many cell culture media supplements (such as N2 and B27 supplements), insulin was present to act upon RGCs in these experiments. Thus, we investigated whether insulin deprivation can lead to a faster disease phenotype in OPTN(E50K) RGCs through a reduced activity of the mTOR signaling pathway. RGCs were grown with medium supplemented with either B27 or B27 without insulin, and RGC neurites were measured from 1 to 4 weeks of maturation following purification (supplemental figure 7A-C). As soon as 2 weeks following purification, OPTN(E50K) RGCs subjected to insulin deprivation exhibited neurite deficits, while the neurites from OPTN(E50K) RGCs with insulin revealed robust outgrowth comparable to wild-type RGCs with or without insulin (Figure 6A-D), as measured by soma size, neurite length, number of primary neurites and Sholl analysis (Figure 6E-H). Insulin deprivation also decreased the mTORC1 effector pS6^Ser240/244^ in OPTN(E50K) RGCs (Figure 6I-K), as well as significantly increased the level of LC3-II (Figure 6L-N), suggesting an imbalance of autophagy and mTORC1 in OPTN(E50K) RGCs when deprived of insulin, resulting in RGC neurite morphological deficits. After a full 4 weeks of RGC growth, consistent with our previous findings, both OPTN(E50K) RGCs with or without insulin exhibited neurite shortening (supplemental figure 7). Our results support the idea that insulin signaling is essential to promote overall RGC neurite outgrowth, and that lack of sufficient mTOR signaling results in neurite retraction.

**Figure 6.**
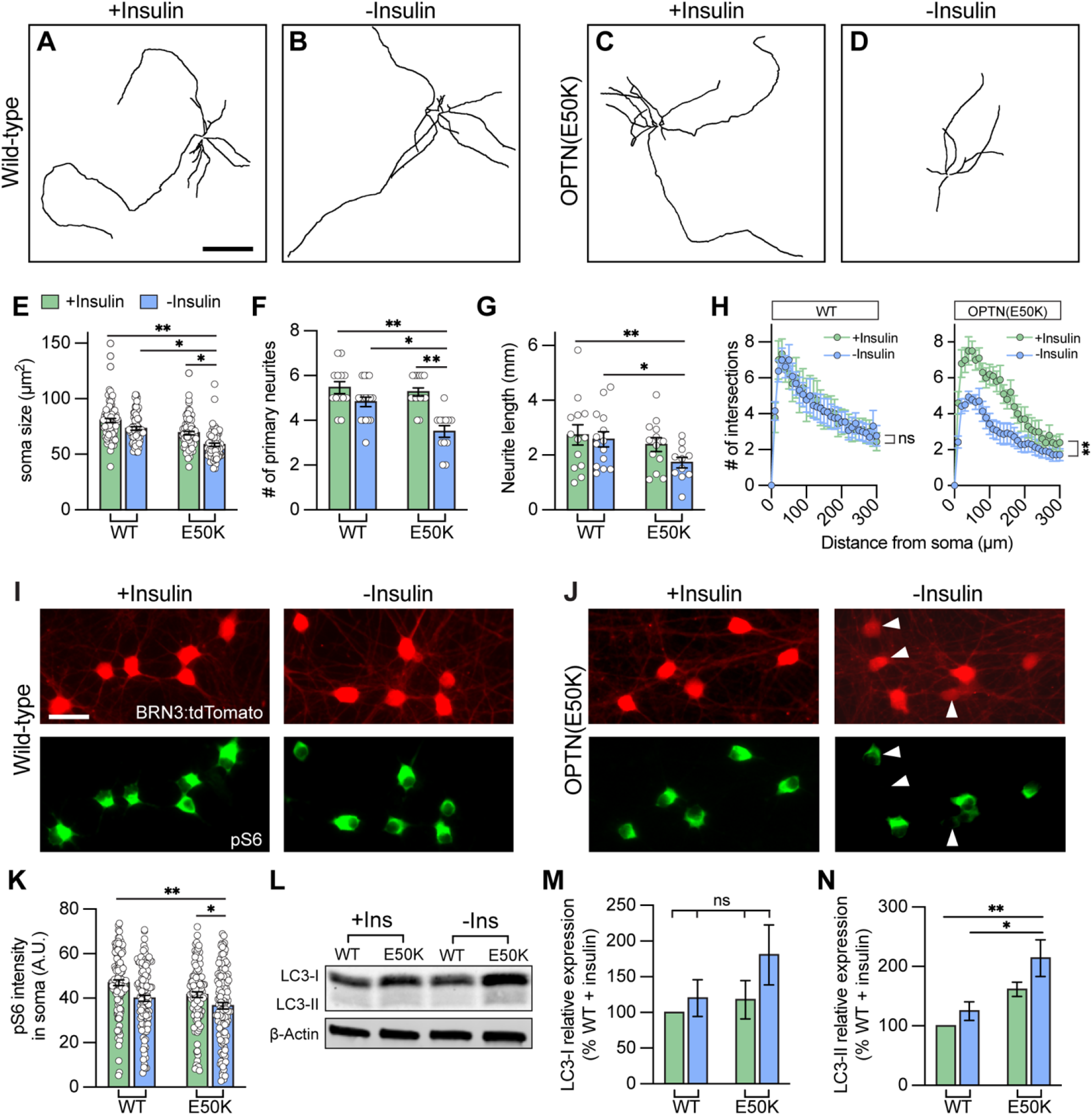
Insulin deprivation accelerated hPSC-RGCs neurite deficits under OPTN(E50K) mutation. (A-D) Representative neurite tracings of WT and OPTN(E50K) hPSC-RGCs after 2 weeks of growth either with insulin or without insulin. Scale bar: 200 μm. (E-H) Quantitative analysis of neurite parameters identified neurite deficits in OPTN(E50K) hPSC-RGCs after insulin deprivation for 2 weeks, as measured by their soma size (n=4 biological replicates using WT(+Ins) n=103, E50K(+Ins) n=91, WT(-Ins) n=74, and E50K(-Ins) n=80 technical replicates; One-way ANOVA, Tukey post hoc test. **p<0.01, *p<0.05) (E), number of primary neurites (n=4 biological replicates using WT(+Ins) n=13, E50K(+Ins) n=15, WT(-Ins) n=17, and E50K(-Ins) n=12 technical replicates; One-way ANOVA, Tukey post hoc test. **p<0.01, *p<0.05) (F), total neurite length (n=4 biological replicates using WT(+Ins) n=13, E50K(+Ins) n=13, WT(-Ins) n=14, and E50K(-Ins) n=11 technical replicates; One-way ANOVA, Tukey post hoc test. **p<0.01, *p<0.05) (G), and Sholl analysis (n=3 biological replicates using WT(+Ins) n=9, E50K(+Ins) n=10, WT(-Ins) n=13, and E50K(-Ins) n=14 technical replicates; t-test, WT(+Ins) vs WT(-Ins): ns= not significant, p=0.73; E50K(+Ins) vs E50K(-Ins): ***p=0.0004) (H). (I-J) Immunostaining revealed that a subset of OPTN(E50K) hPSC-RGCs reduced their expression of pS6^Ser240/244^ when cultured under insulin deprivation, while wild-type RGCs did not change pS6^Ser240/244^ expression under these experimental parameters. Scale bar: 25 μm. (K) Quantification of pS6^Ser240/244^ intensity indicated the decrease of mTORC1 levels in OPTN(E50K) hPSC-RGCs after insulin deprivation (n=3 biological replicates using WT(+Ins) n=119, E50K(+Ins) n=117, WT(-Ins) n=118, and E50K(-Ins) n=126 technical replicates; One-way ANOVA, Tukey post hoc test. **p<0.01, *p<0.05). (L-N), Western blot and quantification of the relative protein expression demonstrated the increased expression of LC3-II in OPTN(E50K) hPSC-RGCs following insulin deprivation for only two weeks (n=5 for each group; One-way ANOVA, Tukey post hoc test. **p<0.01, *p<0.05, ns= not significant, p>0.05). Data are all represented as mean values ± S.E.M. **Figure 6-source data 1. The expression of LC3 protein in hPSC-RGCs with or without insulin deprivation**.

### Trehalose rescues neurodegenerative phenotypes in OPTN(E50K) RGCs by inducing autophagy in an mTOR-independent manner

We have previously identified that autophagy deficits and the accumulation of autophagosomes can be cleared after a short term (24 hr) treatment with the autophagy inducer rapamycin in OPTN(E50K) retinal organoids (VanderWall *et al*., 2020). However, rapamycin induces autophagy via mTOR inhibition, and a reduction in mTOR activity abrogates dendritic outgrowth and maturation in RGCs (Belforte *et al*., 2021), indicating that long term treatment with rapamycin is not an ideal approach. Because our data and others suggest that RGC survival relies upon the homeostatic balance between mTOR and autophagy signaling (Madrakhimov et al., 2021), we hypothesized that inducing autophagy in an mTOR-independent manner can rescue neurodegenerative phenotypes in OPTN(E50K) RGCs through sustained mTOR signaling along with an induction in autophagy. To accomplish this, we used the mTOR-independent autophagy inducer trehalose (25 mM) to stimulate the autophagic-lysosome degradation pathway in RGCs. Following 2 weeks of trehalose treatment, OPTN(E50K) RGCs demonstrated a robust protection of neurite morphology, measured by a preservation of soma size, neurite length, number of primary neurites, and Sholl analysis when compared with wild type RGCs as well as untreated OPTN(E50K) RGCs (Figure 7A-H). Interestingly, 2 weeks of trehalose treatment partially reduced OPTN puncta accumulation in OPTN(E50K) RGCs (Figure 7I-P), while no difference in p62 puncta was observed (supplemental figure 8). Moreover, trehalose treatment reduced LC3-II expression as well as restored levels of the mTORC1 effector pS6^Ser240/244^ (Figure 7Q-T). Collectively, these findings demonstrate that treatment with trehalose can induce autophagy and clear accumulated puncta, while maintaining proper mTOR signaling, leading to sustained overall health of OPTN(E50K) RGCs comparable to their wild type counterparts.

**Figure 7.**
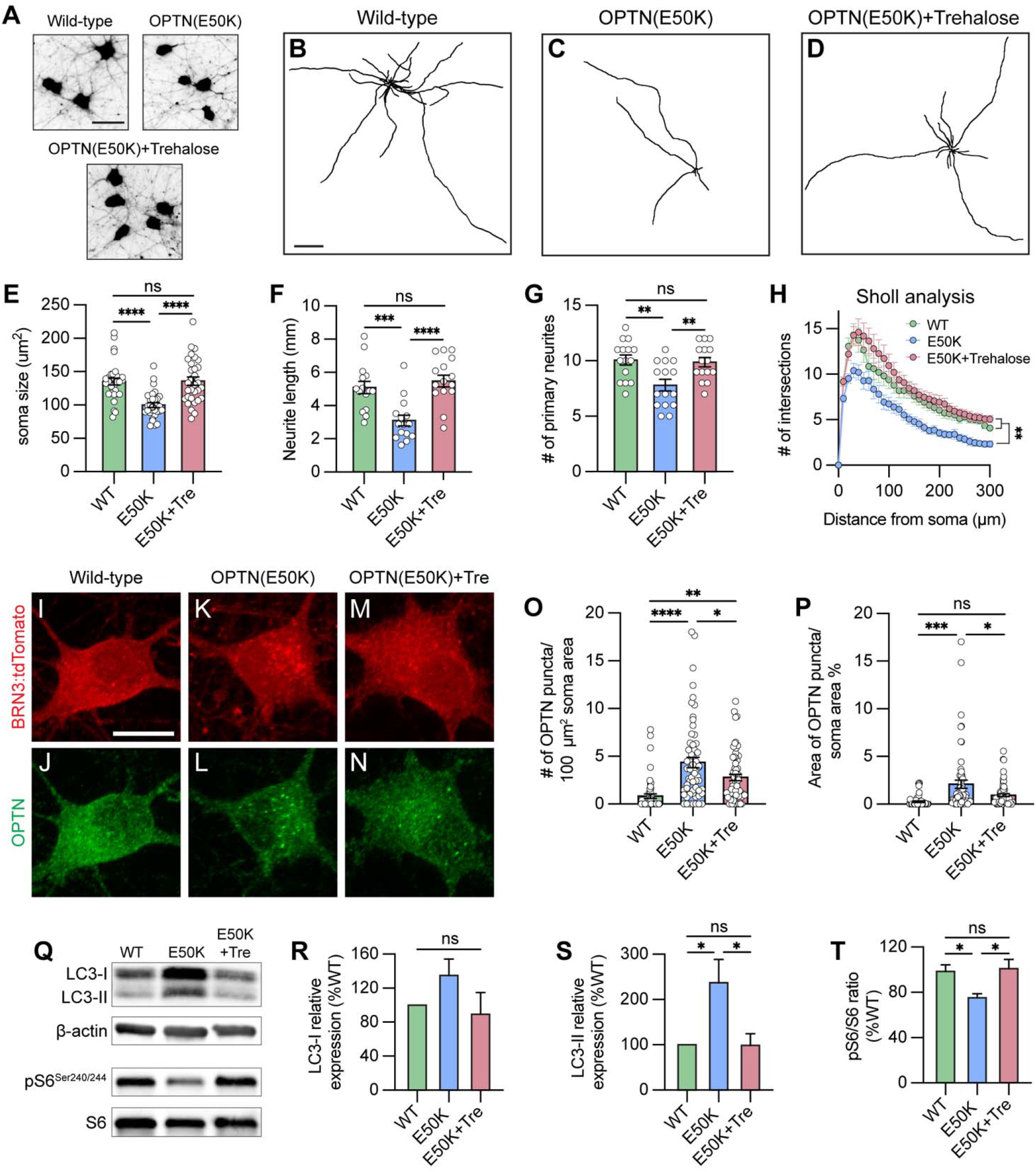
Trehalose prevented neurite retraction and cleared protein accumulation in OPTN(E50K) hPSC-RGCs. (A) Representative images of soma size in wild-type, OPTN(E50K), and trehalose-treated OPTN(E50K) hPSC-RGCs. Scale bar 25 μm. (B-D) Representative neurite tracing images of wild-type, OPTN(E50K), and trehalose-treated OPTN(E50K) hPSC-RGCs. Scale bar: 200 μm. (E-H) Quantitative analysis of neurite parameters demonstrated protection of neurites in OPTN(E50K) hPSC-RGCs after trehalose treatment for 2 weeks, as measured by the soma size (n=3 biological replicates using WT n=30, E50K n=30, and E50K-trehalose n=30 technical replicates, One-way ANOVA, Tukey post hoc test. ****p<0.0001, ns= not significant, p>0.05) (E), number of primary neurites (n=3 biological replicates using WT n=15, E50K n=15, and E50K-trehalose n=15 technical replicates; One-way ANOVA, Tukey post hoc test. ****p<0.0001,***p<0.001, ns= not significant, p>0.05) (F), total neurite length (n=3 biological replicates using WT n=15, E50K n=15, and E50K-trehalose n=15 technical replicates; One-way ANOVA, Tukey post hoc test. **p<0.01, ns= not significant, p>0.05) (G), and Sholl analysis (n=3 biological replicates using WT n=15, E50K n=15, and E50K-trehalose n=15 technical replicates; One-way ANOVA, Tukey post hoc test. WT vs E50K: **p=0.0095; WT vs E50K(trehalose): ns= not significant, p=0.81; E50K vs E50K(trehalose): **p=0.0012) (H). (I-N) Immunostaining displayed the decrease of OPTN puncta in OPTN(E50K) hPSC-RGCs after trehalose treatment. Scale bar: 10 μm. (O-P) Quantification of OPTN puncta in hPSC-RGCs (n=3 biological replicates using WT n=51, E50K n=60, and E50K-trehalose n=61 technical replicates; One-way ANOVA, Tukey post hoc test. ****p<0.0001, ***p<0.001, **p<0.01, *p<0.05, ns= not significant, p>0.05). (Q-T) Western blot and the relative protein expression demonstrated the recovery of changes to LC3-II and pS6^Ser240/244^ expression in OPTN(E50K) hPSC-RGCs after trehalose treatment. (n=5 for each WT, E50K, and E50K with trehalose treatment; One-way ANOVA, Tukey post hoc test. LC3-I: WT vs E50K: p=0.41, WT vs E50K(trehalose): p=0.91, E50K vs E50K(trehalose): p=0.23. LC3-II: WT vs E50K: p=0.03, WT vs E50K(trehalose): p=0.99, E50K vs E50K(trehalose): p=0.03. pS6: WT vs E50K: p=0.048, WT vs E50K(trehalose): p=0.95, E50K vs E50K(trehalose): p=0.029.). Data are all represented as mean values ± S.E.M. **Figure 7-source data 1. The level of LC3 and pS6** ^**Ser240/244**^ **alternation in OPTN(E50K) hPSC-RGCs under trehalose treatment**.

## Discussion

The process of autophagy serves as a cellular protective mechanism by clearing damaged proteins, mitochondria, and organelles within the cells. Ineffective clearance of aggregated proteins in neurons contributes to neurodegenerative diseases such as Alzheimer’s disease, Parkinson’s disease, ALS, and Huntington’s disease (Levine and Kroemer, 2008; Singh et al., 2012). Likewise, recent evidence has demonstrated that dysfunction of the autophagy degradation pathway may be involved in RGC-associated neurodegenerative diseases, including glaucoma, optic atrophy, and diabetic retinopathy (Hirt *et al*., 2018; Madrakhimov *et al*., 2021; White et al., 2009). Glaucoma is characterized by the progressive loss of RGCs, the sole type of projection neuron that connects the retina to the brain. In addition to age, elevated IOP is a prevalent risk factor for glaucoma, while a subpopulation of patients develop glaucoma due to monogenic mutations. Importantly, three monogenic risk genes have been identified in primary open-angle glaucoma (POAG) patients: *MYOC, OPTN*, and *TBK1* (Ritch *et al*., 2014). While these genes account for approximately 5% of POAG patients(Qassim and Siggs, 2020), two of them (*OPTN* and *TBK1*) play prominent roles in autophagy. Indeed, mutations in *OPTN* or *TBK1* can lead to glaucomatous neurodegeneration without elevated IOP, indicating that autophagy dysfunction could contribute to RGC degeneration. Here, we first examined how the OPTN(E50K) mutation contributes to autophagy deficits, resulting in RGC neurodegeneration. We used hPSC-derived RGCs because we previously showed that OPTN(E50K) hPSC-RGCs exhibit morphological and functional deficits compared to isogenic controls, supporting the idea that these cells can serve as an appropriate model to explore RGC neurodegenerative events (VanderWall *et al*., 2020). We identified that this mutation resulted in autophagy disruption based upon increased levels of LC3-II as well as changes in autophagic flux in RGCs with the OPTN(E50K) mutation. Through the use of an LC3-RFP-GFP sensor, we noted a decreased efficiency in autophagosome and lysosome fusion, suggesting that defective autophagy leads to an inability to clear protein accumulation, resulting in an elevation of metabolic stress in RGCs. Significantly, we also explored whether changes in autophagy could be observed more broadly in a magnetic microbead occlusion mouse model. Interestingly, similar to our observations in OPTN(E50K) hPSC-RGCs, we also observed increased LC3-II and changes to autophagic flux in ocular hypertensive retinas, as well as increased levels of LC3 and LAMP1 specifically within RGCs when compared to sham-operated controls. Collectively, our findings support the idea that autophagy dysfunction may be broadly applicable across multiple stressors associated with glaucoma, and highlight the concept that the maintenance of autophagy homeostasis is critical for RGC health by employing two species (human and mouse) as well as different risk factors (gene mutation and intraocular pressure) in glaucoma.

During the process of autophagy, OPTN interacts with LC3 to drive autophagosome maturation(Wong and Holzbaur, 2014). Multiple studies have suggested that changes to OPTN are associated with the appearance of inclusion bodies and promotes neurotoxicity in various neurodegenerative diseases (Anborgh et al., 2005; De Marco et al., 2006; Maruyama et al., 2010; Mori et al., 2012; Osawa et al., 2011; Schwab et al., 2012; Shen et al., 2015; Sirohi and Swarup, 2016). On the contrary, the knockout of OPTN leads to deficits in ubiquitin binding or immune failure, resulting in neural damage (Markovinovic et al., 2018; Munitic et al., 2013; Slowicka et al., 2016). While mutations of OPTN in the LC3-interacting region (LIR) as well as the ubiquitin binding domain (UBD) are known to alter OPTN-associated autophagy (Qiu et al., 2021; Swarup and Sayyad, 2018), the E50K mutation is neither located in the LIR nor UBD.

Enhanced OPTN binding to TBK1 and increased production of insoluble OPTN has been previously observed in E50K mutant cells, suggesting that a loss of OPTN function or a gain of toxic function likely contributes to disease pathogenesis (Li et al., 2016; Minegishi et al., 2013). In animal models, transgenic mice that express the human OPTN(E50K) transgene exhibited RGC loss and axonal degeneration (Tseng et al., 2015), while another study using transgenic mice with OPTN(E50K) overexpression revealed an increased degradation of mitochondria characterized by greater induced mitophagy, resulting in RGC loss (Shim et al., 2016). While OPTN(E50K) genetically modified cell lines or animals do share certain disease-responding mechanisms with human patients, the presence of the wild-type OPTN gene in the genome cannot be excluded in those models. More so, increased levels of OPTN, with or without the E50K mutation, can lead to toxic effects suggesting dose-dependent effects (Park et al., 2006). As a result, the hPSC-RGCs generated through CRISPR/Cas9 gene editing strategies used here are likely a more suitable model to study the functional effects of the OPTN(E50K) mutation in RGCs, mimicking the cellular events that are comparable to patient RGCs that harbor the OPTN(E50K) mutation. To study whether the E50K mutation alters OPTN function, we first characterized its protein expression and localization. Intriguingly, we identified a decreased level of OPTN protein in RGCs with the E50K mutation, in contrast to other studies that overexpressed the mutant protein, suggesting that endogenous OPTN(E50K) protein may be degraded by the ubiquitin-proteosomal system. This reduction of OPTN protein also seems to attenuate the recruitment of LC3 during autophagy, suggesting a loss of protein function in RGCs with the OPTN(E50K) mutation. However, whether the decreased recruitment of LC3 by OPTN directly contributes to defects in the autophagy pathway, or if this recruitment of LC3 can be compensated for by other autophagy receptors is still undetermined (Shoemaker et al., 2019).

Functional deficits in RGCs due to the OPTN(E50K) mutation can be due to either a loss of protein function, or by the mutated protein’s gain of a toxic function through the accumulation of OPTN(E50K) protein aggregates. While we observed that the total level of OPTN protein was decreased in RGCs with the E50K mutation, an accumulation of OPTN aggregates was found in OPTN(E50K) RGCs, indicating that residual OPTN(E50K) protein can still lead to protein accumulation that may be toxic to RGCs. Importantly, our results comparing RGCs to cortical neurons suggests that the accumulation of OPTN(E50K) protein was unique to RGCs, which may then contribute to selective degeneration of RGCs, despite the fact that OPTN plays a role in autophagy within many cell types. Furthermore, by comparing wild-type hPSC-derived neurons, we also observed a higher level of LC3-I in RGCs than in cortical neurons, suggesting that RGCs seem to have a higher demand for autophagosome formation, and that RGCs with the OPTN(E50K) mutation cannot satisfy this requirement, leading to neurodegeneration (Figure 3J and L). It is also important to note that the findings of the current study did not compare RGCs to other types of projection neurons, such as motor neurons. Certain missense mutations in OPTN can cause glaucoma in RGCs, whereas other mutations in OPTN, including deletions, missense, or nonsense mutations, lead to motor neuron loss in ALS (Swarup and Sayyad, 2018). The glaucoma associated mutations do not overlap with ALS associated mutations, and why these mutations selectively promote either RGC or motor neuron degeneration remains unknown. We speculate that one potential reason that the E50K mutation selectively promotes RGC loss can be linked to mitophagy. Since OPTN acts as a receptor particularly for mitophagy, a form of autophagy that selectively degrades unnecessary mitochondria, human RGCs rely heavily on mitochondria for energy supply in the optic nerve head (ONH) as well as the proximal axonal regions prior to the ONH, where RGC axons remain unmyelinated (Wareham et al., 2022). Therefore, a decreased efficiency in the removal of damaged mitochondria could induce metabolic stress, leading to downregulation of metabolic pathways such as mTOR signaling in OPTN(E50K) RGCs.

Our data show that autophagy-induced stress results in changes to the upstream regulator of autophagy, mTORC1. mTORC1 serves as a central nutrient sensor that controls protein synthesis as well as organelle degradation through autophagy during development as well as aging(Kim et al., 2002). mTOR inhibition has been shown to only minimally activate autophagy in primary hippocampal neurons, and tau phosphorylation and Aβ metabolism likely hyperactivate mTOR in Alzheimer’s disease and Down syndrome, indicating that disruption of autophagy in Alzheimer’s disease -related neurodegenerations is mTOR-independent (Linda et al., 2022; Maday and Holzbaur, 2016; Querfurth and Lee, 2021). On the contrary, in RGCs, we and others have demonstrated that mTOR inactivation not only regulated autophagy, but also induced neurite shortening during RGC injury, while additional mTOR stimulation can trigger RGC dendrite and axon regeneration (Agostinone *et al*., 2018; Belforte *et al*., 2021; Duan *et al*., 2015; Madrakhimov *et al*., 2021; Park *et al*., 2008; Teotia et al., 2019), suggesting that mTOR activation is essential for RGC survival. Our previous findings demonstrated neurite retraction in OPTN(E50K) hPSC-RGCs, an indication of protein synthesis attenuation by reduced mTOR signaling, as well as downregulation of mTOR associated pathways from our prior RNA-seq data (VanderWall *et al*., 2020), suggesting that the OPTN(E50K) mutation altered mTOR activity in RGCs. To further confirm this finding, we assessed mTOR effectors and found that OPTN(E50K) hPSC-RGCs selectively downregulated mTORC1 and upregulated AMPK, another nutrient sensor that is activated by energy stress and serves as a negative regulator of mTORC1. However, whether AMPK regulates autophagy directly or indirectly through suppression of mTORC1 still needs to be determined (Inoki et al., 2012; Ji et al., 2015). Interestingly, the hypoactivation of mTORC1 and upregulation of AMPK was also identified in the magnetic microbead occlusion mouse glaucoma model (Belforte *et al*., 2021), further supporting the idea that RGC viability is mTOR-dependent.

Rapamycin and other mTOR antagonists are well-known autophagy inducers that promote clearance of protein aggregates through the inhibition of the mTOR signaling pathway, thereby removing the inhibition upon autophagy (Guertin and Sabatini, 2009). We and others have demonstrated that short term rapamycin treatment is able to clear accumulated autophagosomes and prevent cells from degenerating (Chalasani et al., 2014; VanderWall *et al*., 2020). Nevertheless, since mTOR plays a pivotal role in RGC function as discussed above, long-term exposure to rapamycin may induce adverse effects such as axonal and dendritic degeneration. In addition, it was critical to identify other means to induce autophagy to degrade OPTN(E50K) protein accumulation that did not adversely affect mTOR signaling. To this end, we choose trehalose as a means to modulate autophagy through mTOR-independent means, as trehalose is thought to induce Transcription Factor EB (TFEB) nuclear translocation and activation autophagy-associated proteins independent to mTOR (Palmieri et al., 2017; Rusmini et al., 2019). We observed that trehalose improved autophagy deficits and also rescued neurite retraction in OPTN(E50K) hPSC-RGCs, supporting the idea that mTOR-independent autophagy induction could be a therapeutic target for RGC neurodegeneration by inducing autophagy while maintaining mTOR signaling to promote RGC survival. Taken together, the results of our studies demonstrate a strong connection between autophagy disruption and mTORC1 inactivation, which contributes to neurodegeneration in glaucoma.

## Materials and methods

### Human pluripotent stem cell culture and CRISPR/Cas9 gene editing

For all studies, we used both the H7 human embryonic stem cell line (WiCell Research Institute, RRID: CVCL_9772) as well as a human patient-derived induced-pluripotent stem cell (iPS) cell line with the OPTN(E50K) mutation. Both of these were previously subjected to gene editing using CRISPR/Cas9 techniques to either introduce the OPTN(E50K) mutation or correct the mutation, respectively, as previously described in Vanderwall and Huang et al (VanderWall *et al*., 2020). Additionally, for some experiments, hPSC lines used were previously edited to express a BRN3B-tdTomato-Thy1.2 reporter (VanderWall *et al*., 2020), based of strategies developed by Sluch et al (Sluch et al., 2017). Conversely, for experiments related to the use of the LC3-RFP-GFP sensor in which the tdTomato reporter would interfere with analyses, isogenic pairs of cells were edited that lacked the BRN3b-tdTomato-Thy1.2 reporter following methods previously described(VanderWall *et al*., 2020), and these cell lines were further verified by Sanger sequencing. To maintain all lines of hPSCs, cells were grown on a Matrigel (Corning, cat. no. 354277)-coated plate with mTeSR1 medium (StemCell Technologies, cat. no. 85850). hPSCs were passaged with dispase (2 mg/mL, Life Technologies, cat. no. 17105041) when they reached approximately 70%-80% confluency, every 5-7 days.

### Differentiation of human RGCs

Human retinal organoids were differentiated following previously published methods (Fligor et al., 2020; Meyer et al., 2011; Ohlemacher et al., 2015), and RGCs were subsequently purified and grown in culture to induce further maturation as described previously (VanderWall *et al*., 2020). Briefly, colonies of undifferentiated hPSCs at 80% confluency were enzymatically detached from the plate with dispase and cultured in suspension to induce the formation of embryoid bodies (EBs). EBs were gradually transferred from mTeSR1 to neural induction medium (NIM), which consisted of DMEM/F12(1:1) supplement with N2 supplement, MEM non-essential amino acids and heparin (2 μg/ml). On day 6, BMP4 (50 ng/mL) was added to differentiating cultures, and EBs were subsequently plated on day 8 by supplementation with 10% FBS at a density of approximately 100-200 EBs per well of a 6-well plate to induce a primitive retinal fate. Full medium was changed with NIM on days 9, 12 and 15 and then on day 16, optic vesicle-like structures were lifted and cultured in suspension to induce retinal organoid formation in retinal differentiation medium (RDM), which consisted of DMEM/F12(3:1) supplemented with 2% B27, MEM non-essential amino acids, and penicillin/streptomycin. From day 20-35, half media changes were performed every 2-3 days, and cultures were gradually supplemented with FBS transitioning from 1% to 10%. On day 35, retinal organoids were cultured in retinal maturation medium, consisting of DMEM/F12(3:1) supplement with 2% B27 supplement, MEM non-essential amino acids, penicillin/streptomycin, 10% FBS, 1X GlutaMAX, and 100 μM Taurine. Half media changes were then performed every 2-3 days.

To isolate RGCs, retinal organoids between 45-55 days of differentiation were enzymatically dissociated with AccuMax and purified by Magnetic Activated Cell Sorting (MACS) with CD90.2 (Thy1.2) MicroBeads (Miltenyi Biotec, cat. no. 130-121-278) as previously described (Sluch *et al*., 2017). Purified RGCs were plated on either poly-D-ornithine and laminin-coated glass coverslips or laminin-coated culture plates at a density of 450 cells/mm^2^ and maintained in Brainphys medium supplemented with 2% B27 (either with or without insulin, depending on experiment), 20 ng/mL BDNF, 20 ng/mL GDNF, 1 mM dibutyryl cAMP, 200 nM ascorbic acid and penicillin/streptomycin. Half media changes were performed every 3-4 days and maintained up to an additional 4 weeks. For chloroquine treatment, 30 nM chloroquine (Life Technologies, cat. no. P36239B) was added to hPSC-RGCs after 4 weeks of maturation for a duration of 16 hours prior to fixation and immunohistochemistry. For mTOR inhibition, the dual mTOR inhibitor KU-0063794 (Tocris, cat. no. 3725) was added at indicated concentrations at 3 days following the isolation of RGCs for an additional week. For RGC protection, 25 mM trehalose (MP Biomedicals, cat. no. 103097) was dissolved in the medium, filtered, and added to RGCs from week 2 to week 4 following purification.

### Differentiation of human cortical neurons

To derive cortical neurons from hPSCs, maintenance, passaging, as well as EB formation were performed as described above for retinal organoids with minor modifications (Fligor *et al*., 2020). Primarily, on day 6 of differentiation, 200 nM LDN-193189 (Reprocell, cat. no. 04-0074-02) was added to EBs instead of BMP4. At day 16, the loosely attached rosettes were mechanically lifted and cultured in suspension in RDM as cortical organoids. By day 45, cortical organoids were then enzymatically dissociated with AccuMax and purified by MACS using anti-PSA-NCAM MicroBeads (Miltenyi Biotec, cat. no. 130-097-859) and plated onto poly-D-ornithine and laminin-coated coverslips or laminin-coated culture plates and further maintained in Brainphys medium, as described above.

### Magnetic microbead occlusion mouse glaucoma model

All animal procedures were approved by the University of Montreal Hospital Research Centre (2021-9727, N18025ADPs) and followed the Canadian Council on Animal Care guidelines. Experiments were performed in adult female C57B6L/6 mice (2 to 3 months of age, 20 to 25.1 g) (Charles River, Strain code:027). All procedures were performed under general anesthesia (20 mg/kg ketamine, 2 mg/kg xylazine, 0.4 mg/kg acepromazine). Unilateral elevation of intraocular pressure was performed by a single injection of magnetic microbeads into the mouse anterior chamber(Ito *et al*., 2016). Animals were anesthetized and a drop of tropicamide was applied on the cornea to induce pupil dilation (Mydriacyl, Alcon, Mississauga, ON, Canada). A custom-made sharpened microneedle attached to a microsyringe pump (World Precision Instruments, Sarasota, FL) was loaded with a magnetic microbead solution (1.5 μl, diameter: 4.5 μm, 2.4 × 106 beads, Dynabeads M-450 Epoxy, ThermoFisher Scientific, Waltham, MA). Using a micromanipulator, the tip of the microneedle was gently pushed through the cornea to inject the microbeads into the anterior chamber. The microbeads were immediately attracted to the iridocorneal angle using a hand-held magnet. This procedure avoided injury to ocular structures including the lens and iris. Sham-operated controls received an injection of phosphate buffered saline (PBS). Only one eye was operated on and an antibiotic drop (Tobrex, tobramycin 0.3%, Alcon) was applied immediately after the surgery. The animal was allowed to recover on a heating pad. Intraocular pressure was measured in awake animals before and after the procedure, and bi-weekly thereafter always at the same time using a calibrated TonoLab rebound tonometer (Icare, Vantaa, Finland). For this purpose, a drop of proparacaine hydrochloride (0.5%, Alcon) was applied to the cornea and a minimum of 10 consecutive readings were taken per eye and averaged.

### RNA extraction and quantitative real-time PCR (qPCR)

Total RNA was collected from RGCs following purification and subsequent maturation for 4 weeks using the PicoPure RNA isolation kit (Life Technologies, cat. no. KIT0204). 1 μg of RNA was used for reverse transcription using the High Capacity RNA-to-cDNA kit (Life Technologies, cat. no. 4387406), and cDNA samples were diluted with nuclease-free water at a 1:2 ratio. Quantitative PCR was performed using the Applied Biosystems StepOnePlus Real-Time PCR System with SYBR Green PCR master mix (Life Technologies, cat. no. 4364344) Primer sets used included: OPTN-forward: GACACGTTACAGATTCACGTGA; OPTN-reverse: ACTGTGCCCGGCCTGTTTTC; β-actin-forward: GCGAGAAGATGACCCAGATC; β-actin-reverse: CCAGTGGTACGGCCAGAGG; GAPDH-forward: CGCTCTCTGCTCCTCCTGTT; GAPDH-reverse: CCATGGTGTCTGAGCGATGT. In addition, a melt-curve analysis was performed immediately after the amplification protocol to verify the specificity of amplification. β-actin and GAPDH were used as endogenous controls to normalize the expression levels. Relative mRNA levels were calculated using the ΔΔCT equation (Livak and Schmittgen, 2001).

### Immunoblotting

hPSC-RGC samples were collected in M-PER Mammalian Protein Extraction Reagent (Life Technologies, cat. no. 78501) supplemented with protease and phosphatase inhibitor cocktail (Life Technologies, cat. no. 78440). Alternatively, animals were sacrificed by cervical dislocation and the retinas were immediately dissected out in cold PBS, snap-frozen, and kept at -80°C until protein extraction. Proteins were extracted by homogenization of retinas in lysis buffer (Tris 50 mM, EDTA 1 mM, NaCl 150 mM, NP-40 1% v/v, NaVO3 2mM, NaF 5 mM, Na Deoxycholate 0.25%). The homogenized protein samples were combined with 4x sample buffer and 25μM DTT and incubated at 100°C for 10 minutes followed by loading into a 4-15% gradient pre-cast gel and transferred onto a nitrocellulose membrane through the Trans-Blot Turbo system (BioRad). The membrane was blocked in 5% milk in tris buffered saline (TBS) supplemented with 0.1% Tween-20 (TBS-T) for 30 minutes. The membrane was then incubated with diluted primary antibodies in TBS-T with 5% BSA overnight at 4°C. The membrane was washed 3 times with TBS-T on the following day, and the appropriate secondary antibody in 5% milk in TBS-T was applied for an hour at room temperature, protected from light. The membrane was then washed 3 times with TBS-T and imaged using the Li-COR Odyssey CLx imaging system. Protein intensities were quantified for each band and normalized to an internal control (β-actin) using ImageJ. Detailed information regarding primary antibodies as well as the respective dilutions are provided in the Supplemental Table.

### Immunostaining

Purified hPSC-RGCs were plated on Poly-D-Ornithine and laminin-coated 12 mm coverslips at a density between 20,000 to 50,000 cells. RGCs were fixed at indicated timepoints with 4% paraformaldehyde for 30 min. Alternatively, animals were anesthetized and transcardially perfused with ice-cold 4% paraformaldehyde (PFA) in PBS. Eyes were immediately collected, post-fixed in PFA, and processed to generate cryosections as previously described(Pernet et al., 2007). Briefly, the cornea and the lens were removed, and the eye cup was incubated in the same fixative for 2h at 4°C. Tissue was further incubated in 30% sucrose overnight, and then frozen in optimal cutting temperature (O.C.T.) compound (Tissue-Tek, Miles Laboratories, Elkhart, IN, USA). Retinal cryosections (16μm) were collected onto gelatin-coated slides for immunostaining. hPSC-RGCs and retinal cryosections were permeabilized with 0.1% Triton-X for 10 min at room temperature. Following three washes in PBS, RGCs were then blocked in 10% donkey serum and 1% BSA for 1 hour at room temperature. Primary antibodies (Supplemental Table) were diluted in PBS supplemented with 10% donkey serum and 1% BSA and applied overnight at 4°C. On the following day, RGCs were washed three times with PBS and incubated with secondary antibodies diluted in 10% donkey serum and 1% BSA for 1 hour at room temperature. RGCs were then washed three times with PBS and the coverslips mounted onto slides for imaging. Imaging was performed using either a Nikon A1R Confocal Microscope or a Leica DM5500 fluorescence microscope.

Quantification of OPTN puncta was performed in Fiji where particles were analyzed with proper threshold. For LC3 and OPTN colocalization analysis, the LC3B-GFP Sensor (Life Technologies, cat. no. P36235) was added to hPSC-RGCs for 16 hours. RGCs were then stained with OPTN and RFP antibodies, with the latter used to enhance the tdTomato signal. The colocalization between LC3 and OPTN puncta was analyzed within RGC somas by using Fiji with the JACop plugin with threshold.

### Autophagosome maturation assay

To analyze autophagic flux, hPSCs with the OPTN(E50K) mutation as well as isogenic controls both lacking the BRN3b-tdTomato-Thy1.2 reporter were used. Differentiated retinal organoids were enzymatically dissociated with AccuMax and plated on 12 mm coverslips in Brainphys medium for 4 weeks, as described above. Dissociated cells were treated with 1 μM of 5-fluoro-2′-deoxyuridine (Sigma, cat. no. F0503) for the first 24 hours to remove presumptive progenitor and/or glial cells. The RFP-GFP-LC3B sensor (Life Technologies, cat. no. P36239) was added to hPSC-RGCs after 4 weeks of maturation for a duration of 16 hours.

Subsequently, immunohistochemistry was performed to stain the cells with a BRN3 primary antibody to definitively identify RGCs, followed by an Alexa Fluor 647 anti-goat secondary antibody. Immunofluorescence images were visualized on a Nikon A1R Confocal Microscope with Z-stack. To ensure the proper selection of RGCs, only cells expressing BRN3 were considered for further analysis of RFP-GFP-LC3B expression. Quantification of autophagosomes (both RFP and GFP puncta) and autolysosomes (RFP positive, GFP negative puncta) was performed on RGC somas through Fiji by using JACop plugin with appropriate threshold. Autophagosomes were calculated as the fraction of RFP puncta overlapping with GFP puncta.

### Neurite tracing and analysis

hPSC-RGCs or cortical neurons were maintained on 12 mm coverslips. To identify neurites from individual BRN3-tdTomato (RGCs) or β-III Tubulin (cortical neurons) cells, cultures were transfected either with a GFP plasmid using Lipofectamine Stem Reagent (Life Technologies, cat. no. STEM00003) or BacMam GFP (Life Technologies, cat. no. B10383), which provided a relatively low efficiency of transfection that allowed for facile analysis of individual neurons within otherwise dense neuronal cultures that allowed for greater viability of neurons. Transfection was performed two days before fixation, and immunohistochemistry was then performed on these cultures as indicated. Images were taken using a Leica DM5500 fluorescence microscope. Neurite tracing and Sholl analyses were performed using Fiji and the neuroanatomy plugin.

### Proteomic analysis by mass spectrometry

Four biological replicates of wild-type and OPTN(E50K) hPSC-RGCs were lysed with 250 μl of 8 M urea in 100 mM Tris HCl, pH 8.5. Following Bradford assay for protein quantitation (Protein Assay Dye Reagent Concentrate, Bio-Rad, Cat No: 5000006), 40 μg of each protein sample was reduced with 5 mM tris (2-carboxyethyl) phosphine hydrochloride (TCEP, Sigma-Aldrich, Cat No: C4706) and then diluted in 50 mM Tris HCl. Samples were then digested overnight at 35ºC using Trypsin/Lys-C (Trypsin/Lys-C Mix, Mass Spec Grade, Promega, Cat No: V5072); the enzyme-substrate ratio of 1:70). Digestions were quenched with trifluoracetic acid (TFA, 0.5% v/v) and desalted on Waters Sep-Pak cartridges, followed by elution in 50% and 70% acetonitrile containing 0.1% formic acid (FA), and then dried and stored at -20ºC. Dried peptides were later reconstituted in 50 mM triethylammonium bicarbonate pH 8.0 (TEAB). Peptides were labeled for two hours at room temperature with 0.2 mg of Tandem Mass Tag (TMT) reagent (TMT™ Isobaric Label Reagent Set, Thermo Fisher Scientific, Cat No: 90110, Lot No: WG320953). Labeled peptides were then pooled and dried, and then reconstituted in 0.1% TFA and half was fractionated using a Waters Sep-Pak cartridge (Waters™, Cat No: WAT054955) with a wash of 1 mL 0.1% TFA and 0.5% acetonitrile containing 0.1% triethylamine followed by elution in 10%, 12.5%, 15%, 17.5%, 20%, 22.5%, 25% and 70% acetonitrile containing 0.1% triethylamine. A tenth of each fraction was separated on a 25 cm aurora column (IonOpticks, AUR2-25075C18A) at 400 nL/min in the EASY-nLC HPLC system (SCR: 014993, Thermo Fisher Scientific). Nano-LC-MS/MS data were acquired in Orbitrap Eclipse™ Tribrid mass spectrometer (Thermo Fisher Scientific) with a FAIMS pro interface. The data were recorded using Thermo Fisher Scientific Xcalibur (4.3) software (Thermo Fisher Scientific Inc.).

### Statistical analysis

Data in all experiments is represented as mean ± SEM and n represents the number of technical replicates across all experiments. Statistical comparisons were conducted by either Student’s t-test or ANOVA with Tukey post hoc test (specified in figure legends) using GraphPad Prism 9. Statistically significant differences were defined as p < 0.05 in all experiments including proteomics. For proteomics data analysis, RAW files were analyzed in Proteome Discover™ 2.5 (Thermo Fisher Scientific, RRID: SCR_014477) with a Homo sapiens UniProt reviewed FASTA and common contaminants (20417 total sequences). Normalized abundance values for each sample type, abundance ratio, log2(abundance ratio) values, and respective p-values (t-test) from Proteome Discover™ were exported to Microsoft Excel. All processed and raw data are uploaded to ProteomeXchange Accession PXD033173.

## Acknowledgements

We thank Peter Bor-Chian Lin and Adrian Oblak for sharing reagents, as well as Mallika Valapala for helping discussions regarding trehalose experiments. We also thank Amber Mosley and Emma Doud at the IU School of Medicine Center for Proteome Analysis for assistance with proteomics experiments, and the IU School of Medicine Flow Cytometry Core Facility for assistance with cell sorting. Grant support was provided by the National Eye Institute (R01EY033022 to JSM), the BrightFocus Foundation (G2020369 to JSM), the Glaucoma Research Foundation (to JSM), and the Indiana Department of Health Spinal Cord and Brain Injury Research Fund (26343 to JSM). Support for this project was also provided by the Sarah Roush Memorial Fellowship from the Indiana Alzheimer’s Disease Center (CG). This publication was also made possible with partial support from the Stark Neurosciences Research Institute/Eli Lilly and Co. Neurodegeneration fellowship (KCH), from a pre-doctoral fellowship award (KBV) from the National Institutes of Health, National Center for Advancing Translational Sciences, Clinical and Translational Sciences Award (UL1TR002529, A. Shekhar, PI), and from Sigma Xi Grants in Aid of Research (G03152021117541788 to KCH). This work was also partially funded by a grant to ADP from the National Institutes of Health (NIH, 1R01EY030838-01). YS was supported by postdoctoral fellowships from the Fonds de recherche Santé – Québec (FRQS, 276181) and the Canadian Institutes of Health Research (CIHR, 458569).

## Supporting information

**Supplemental Figure 1.**
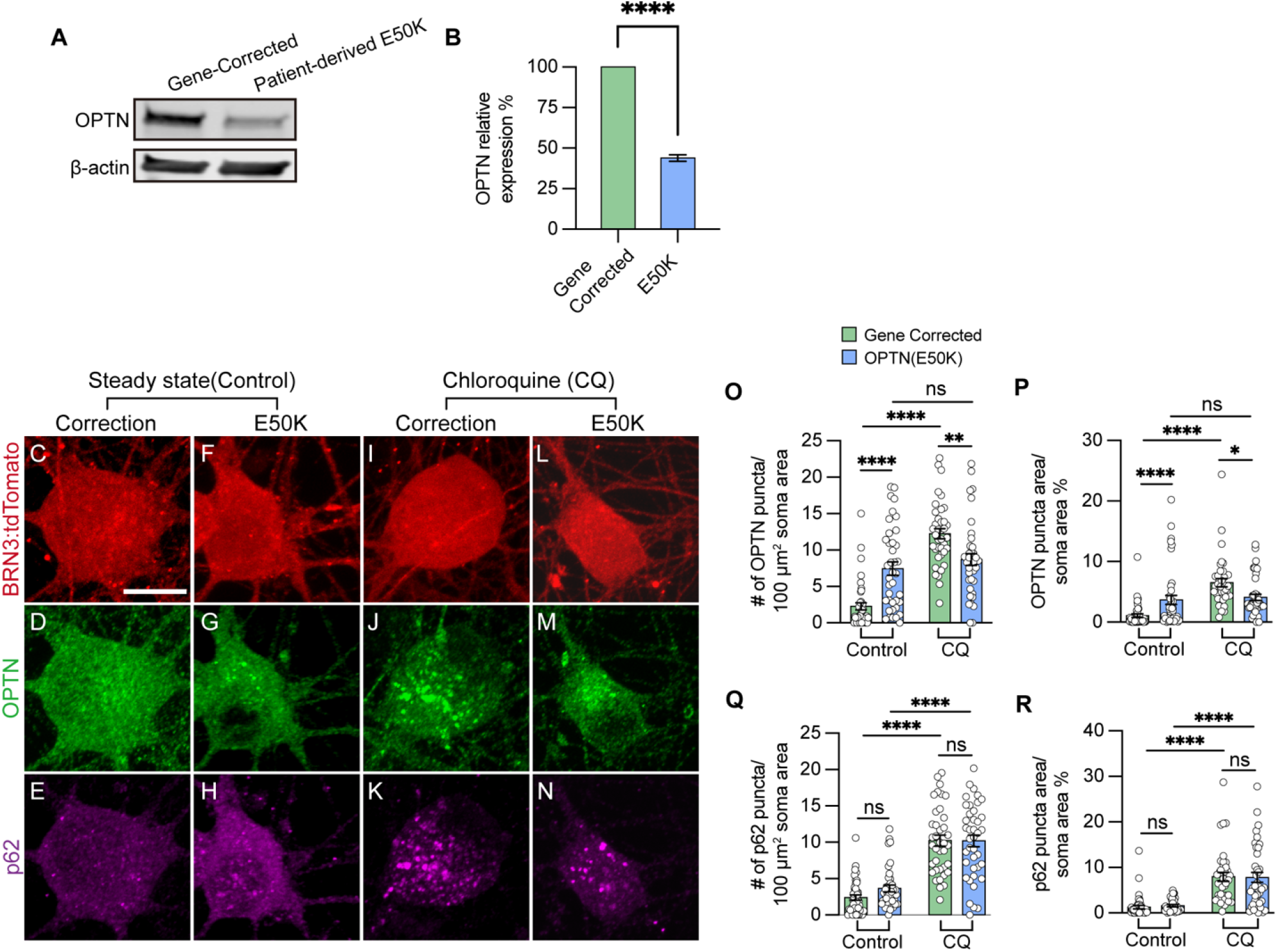
Confirmation of changes in OPTN levels in RGC-derived from patient-specific iPSCs with OPTN(E50K) mutation compared to corresponding isogenic control line. (A-B) Western blot of OPTN relative to β-actin in patient-derived OPTN(E50K) and gene-corrected isogenic control iPSC-RGCs (n=5 for each WT and E50K; t-test, ****p<0.0001). (C-N) Immunostaining revealed the aggregation of OPTN and p62 puncta in BRN3:tdTomato iPSC-RGCs in patient-derived OPTN(E50K) mutant and gene-corrected isogenic control lines after steady state (control) (C-H) and chloroquine (CQ) treatment (I-N). Scale bar: 10 μm. (O-R) Quantification of OPTN puncta (O and P) or p62 puncta (Q and R) in iPSC-RGC (n=3 biological replicates using Ctrl-WT n=42, Ctrl-E50K n=39, CQ-WT n=38 and CQ-E50K n=40 technical replicates; One-way ANOVA, Tukey post hoc test. ****p<0.0001, **p<0.01, *p<0.05, ns= not significant). Data are all represented as mean values ± S.E.M. **Supplemental figure 1-source data 1. The alternation of OPTN protein in gene corrected and patient derived-OPTN(E50K) mutation hPSC-RGCs**.

**Supplemental Figure 2.**
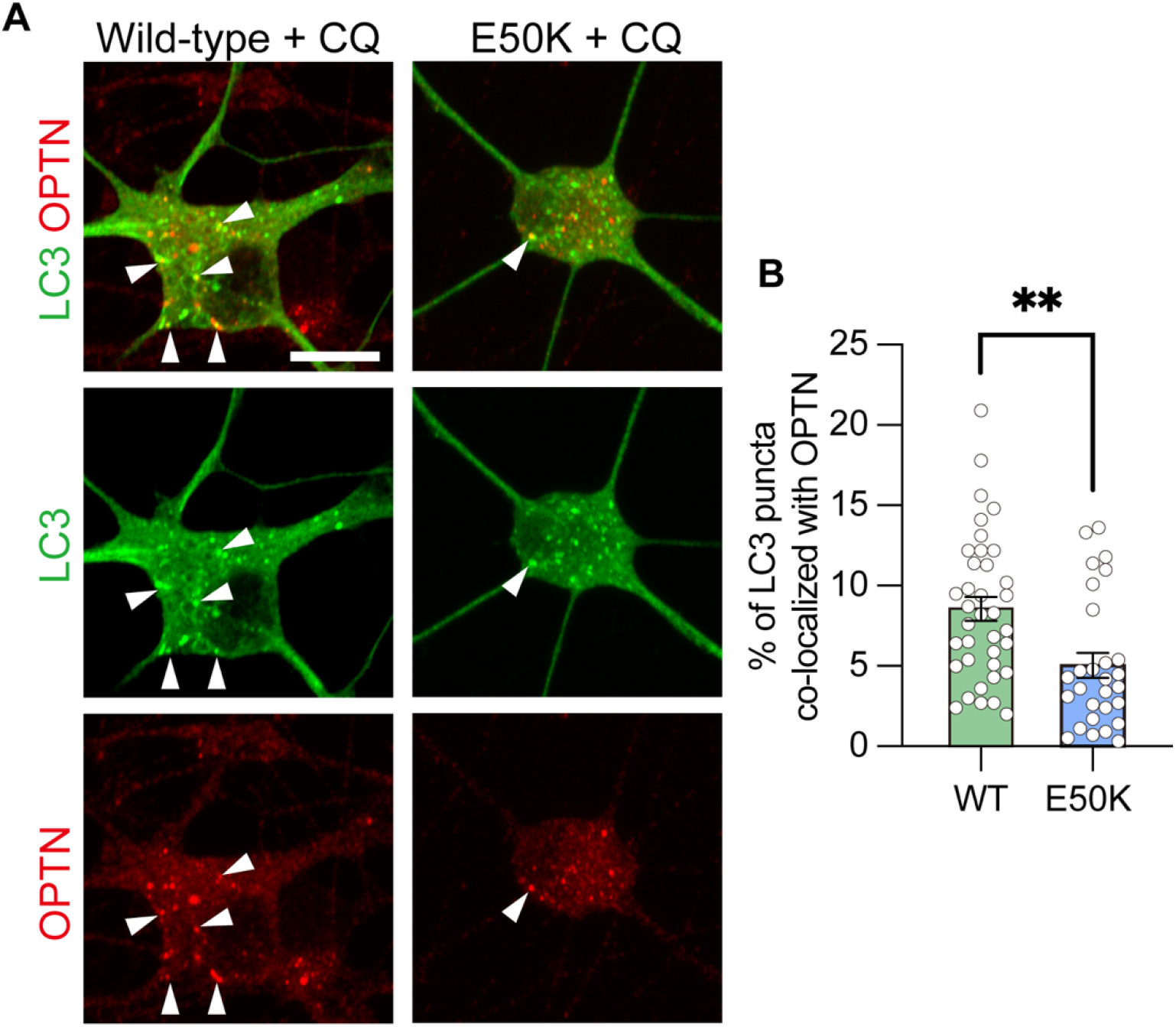
Confirmation of reduced recruitment of LC3 by OPTN in patient-derived OPTN(E50K) iPSC-RGCs after chloroquine treatment. (A) Representative images of OPTN and LC3 localization in patient-derived iPSC-RGCs from wild-type and OPTN(E50K) cell lines after chloroquine treatment. White arrows identify puncta colocalized with OPTN and LC3. (B) Quantification of colocalization between OPTN and LC3 in patient-derived iPSC-RGCs (n=3 biological replicates using WT n=37 and E50K n=28 technical replicates; t-test, **p>0.005). Scale bar: 10 μm. Data are all represented as mean values ± S.E.M.

**Supplemental Figure 3.**
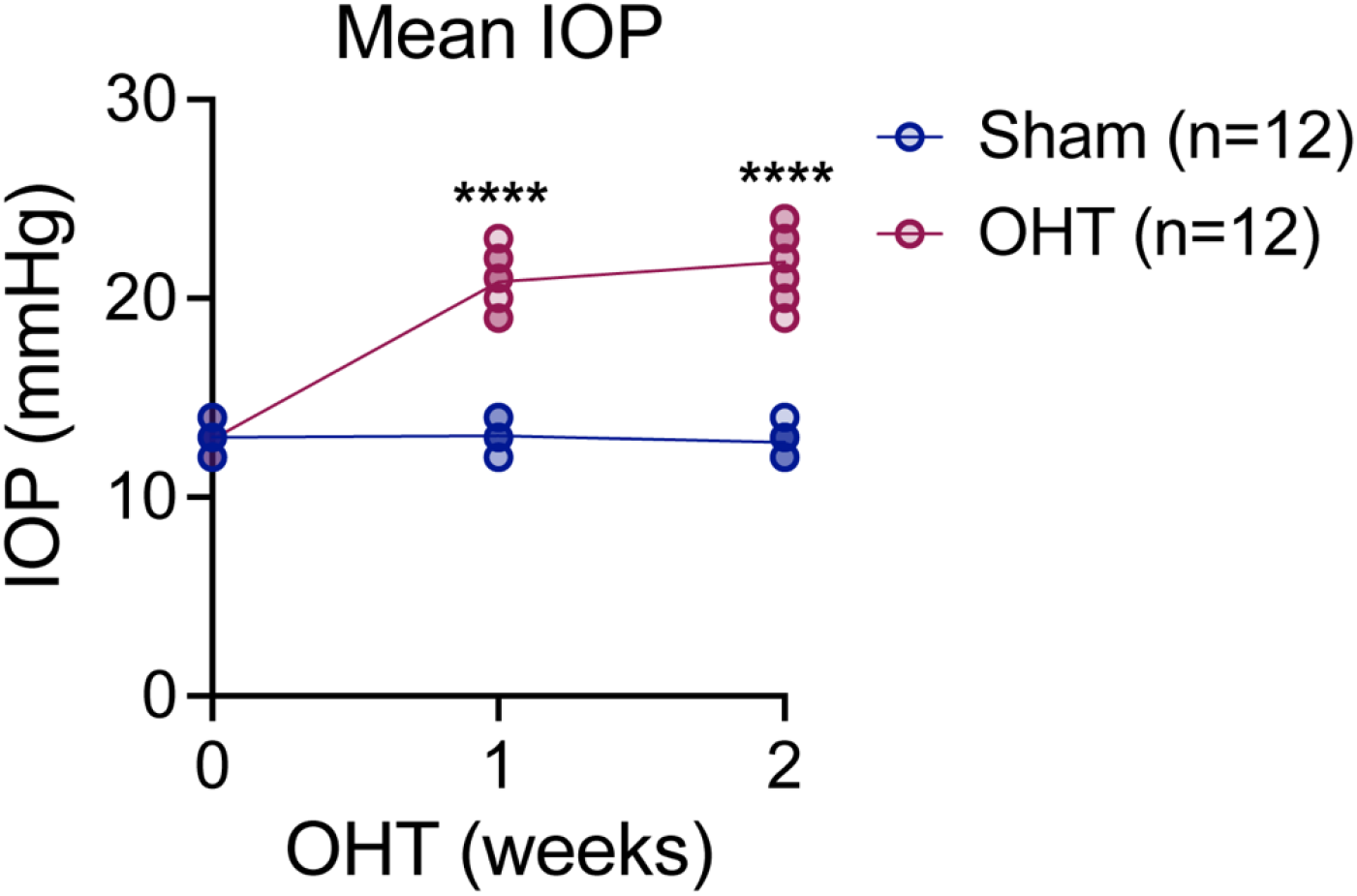
Elevation of intraocular pressure in an ocular hypertension glaucoma model. Maintained elevation of intraocular pressure in mouse eyes following the injection of magnetic microbeads into the mouse anterior chamber, compared to sham-injected controls. Two-way ANOVA with Tukey’s multiple comparison pos hoc test, ****p < 0.0001, n = 12 mice/group.

**Supplemental Figure 4.**
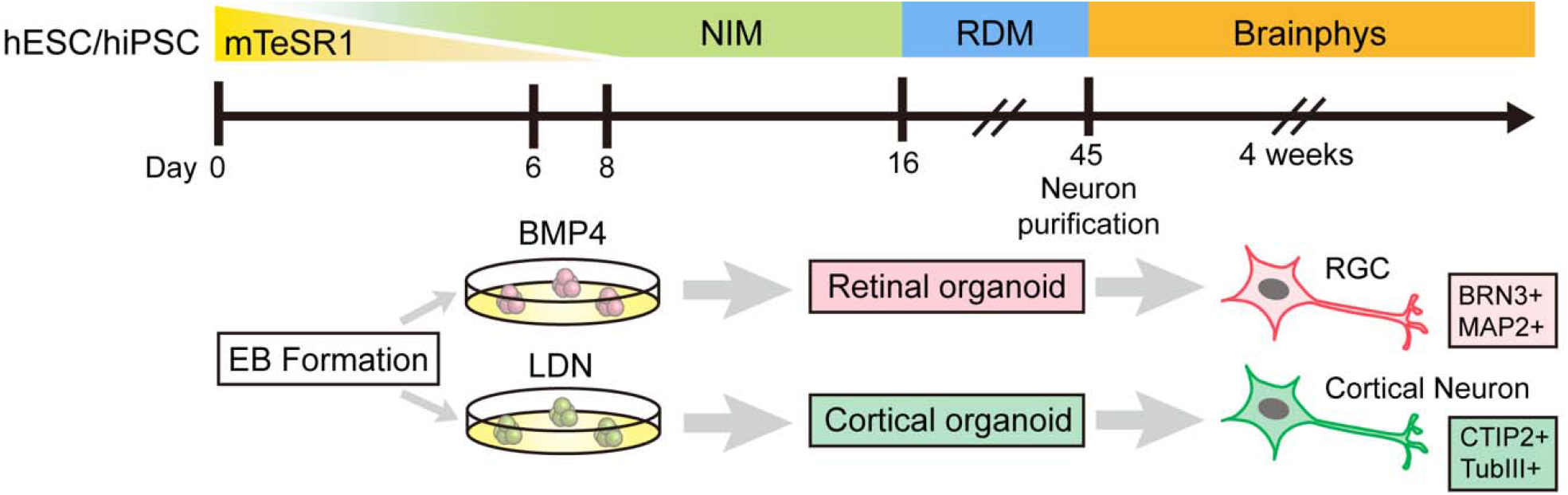
Schematic diagram outlining the differentiation of RGCs and cortical neurons from hPSCs.

**Supplemental Figure 5.**
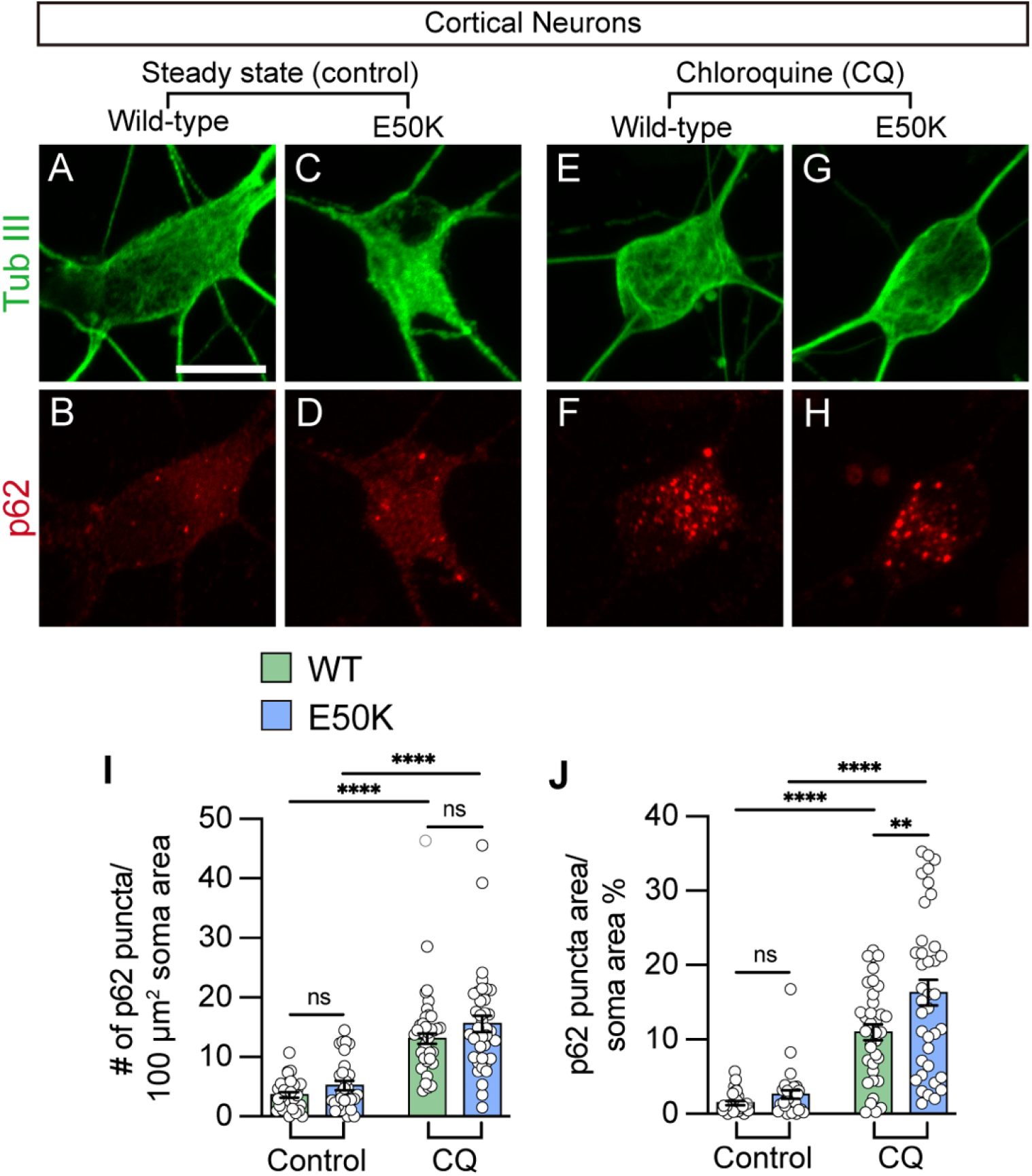
Characterization of p62 expression in hPSC-cortical neurons from wild-type and OPTN(E50K) cell lines. (A-H) Representative images of p62 puncta in iPSC-derived cortical neurons from WT and OPTN(E50K) cell lines under steady state (control) (A-D) and chloroquine (CQ) treatment (E-H). Scale bar: 10 μm. (I-J) Quantification of p62 puncta in iPSC-derived cortical neurons (n=3 biological replicates using Ctrl-WT n=30, Ctrl-E50K n=30, CQ-WT n=36 and CQ-E50K n=38 technical replicates; One-way ANOVA, Tukey post hoc test. ****p<0.0001, **p<0.01, ns= not significant, p>0.05). Data are all represented as mean values ± S.E.M.

**Supplemental Figure 6.**
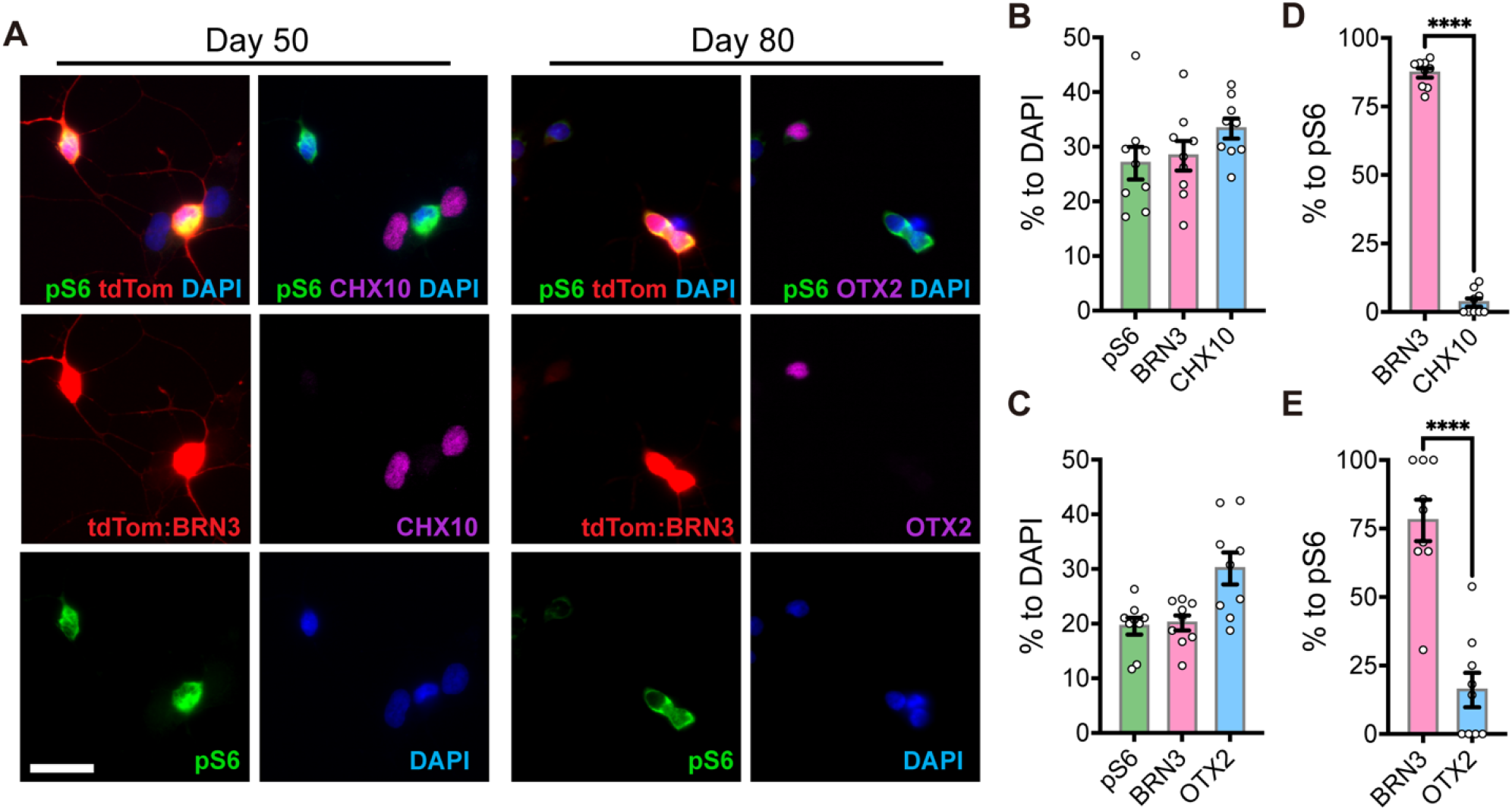
mTORC1 activity is preferentially observed within RGCs among hPSC-derived retinal cells. (A) Retinal organoids were dissociated and plated onto laminin-coated coverslips at either day 50 or day 80 of differentiation to acquire the majority of major retinal cell types, including RGCs (BRN3:tdTomato), retinal progenitor cells (CHX10), and photoreceptors (OTX2), following by analysis of mTORC1 activity based upon co-staining with pS6Ser240/244. Scale bar: 25 μm. (B-C) Quantification of results showing the percentage of retinal cell types observed at day 50 or day 80 (n=9 images from three technical replicates). (D-E) Quantification of pS6Ser240/244 expression colocalized with either BRN3B:tdTomato, CHX10 or OTX2, suggesting that mTORC1 signaling is highly expressed in RGCs, with little expression in retinal progenitor cells or photoreceptors (n=9 images from three technical replicates; t-test, BRN3 vs CHX10: p<0.0001, BRN3 vs OTX2: p<0.0001). Data are all represented as mean values ± S.E.M.

**Supplemental Figure 7.**
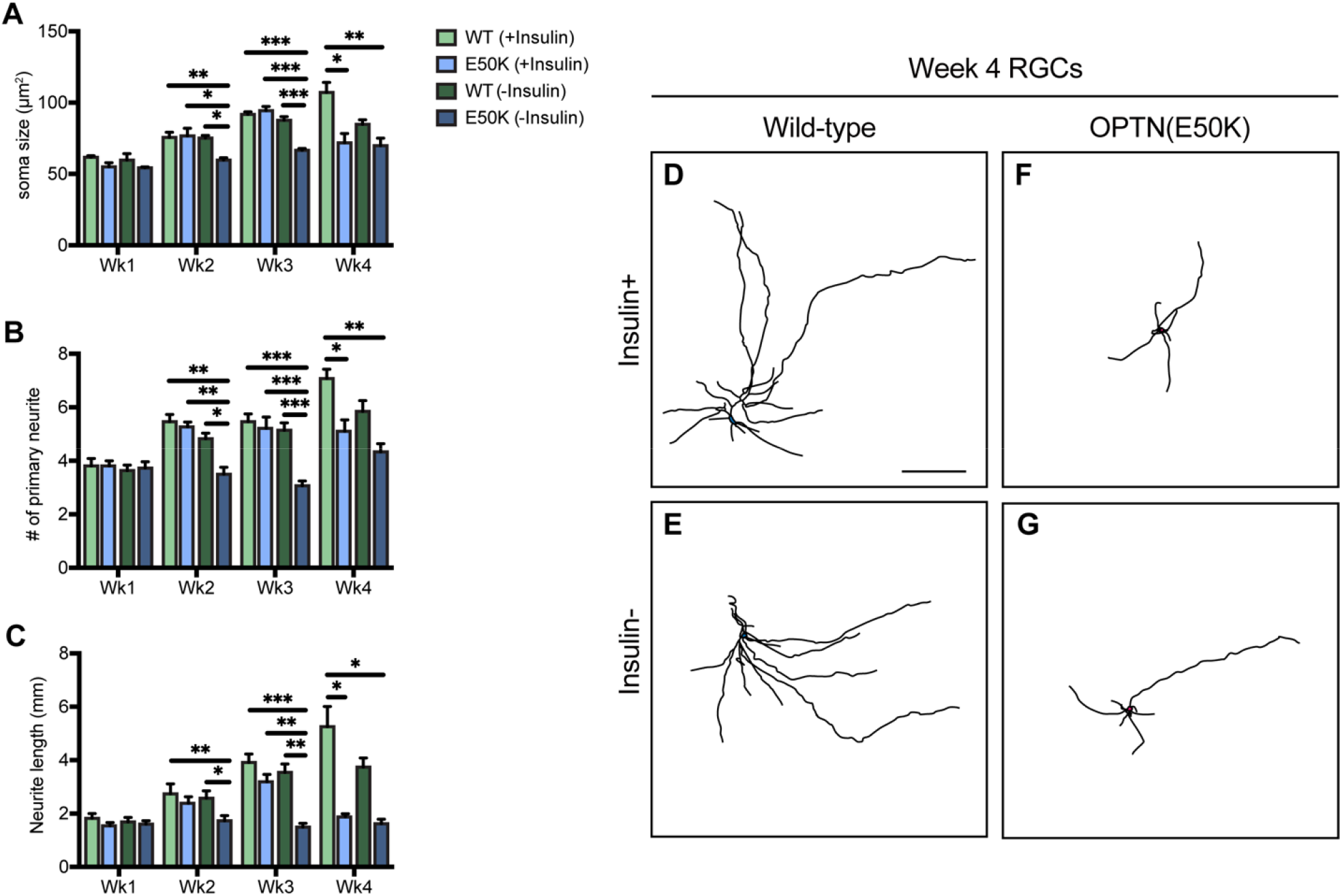
Insulin deprivation expedites the onset of degenerative phenotypes in hPSC-RGCs with the OPTN(E50K) mutation. (A-C) Quantitative analysis of neurite measurements over the course of 4 weeks of differentiation in wild-type and OPTN(E50K) hPSC-RGCs grown with or without insulin, as measured by soma size (n≥30 each condition with 4 biological replicates; One-way ANOVA, Tukey post hoc test. ***p<0.001, **p<0.01, *p<0.05) (A), number of primary neurites (n≥10 each condition with 4 biological replicates; One-way ANOVA, Tukey post hoc test. ***p<0.001, **p<0.01, *p<0.05) (B), total neurite length (n≥10 each condition with 4 biological replicates; One-way ANOVA, Tukey post hoc test. ***p<0.001, **p<0.01, *p<0.05) (C). Data are all represented as mean values ± S.E.M. (D-G) Representative neurite tracings of WT and OPTN(E50K) hPSC-RGCs after 4 weeks of growth either with or without insulin. Scale bar: 200 μm.

**Supplemental Figure 8.**
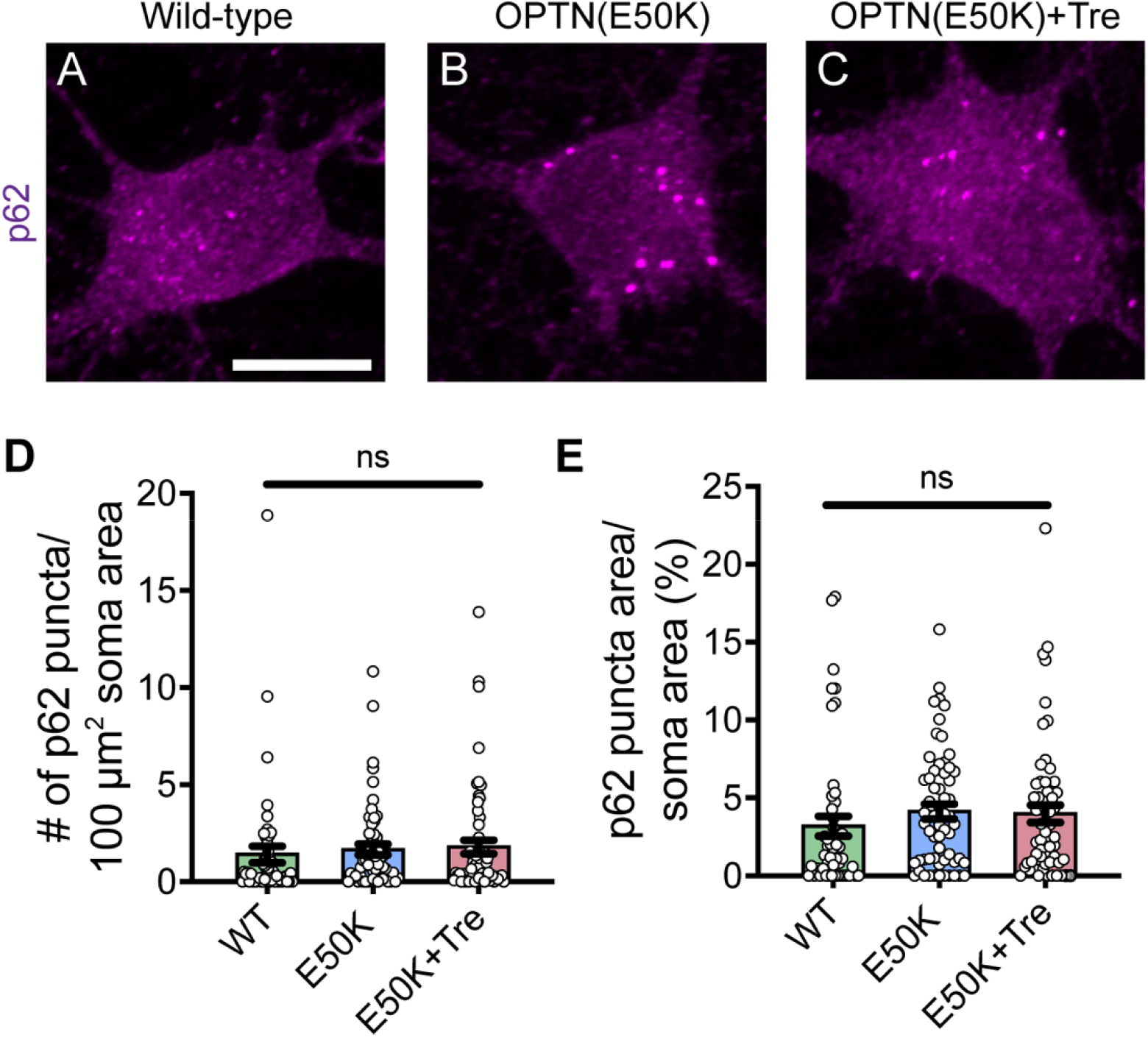
p62 expression remains unchanged in hPSC-RGCs comparing wild-type, OPTN(E50K) and OPTN(E50K) plus trehalose conditions. (A-C) Representative images of p62 puncta in hPSC-RGCs. Scale bar: 10 μm. (D-E) Quantification of p62 puncta in hPSC-RGCs (n=3 biological replicates using WT n=51, E50K n=60, and E50K-trehalose n=61 technical replicates; One-way ANOVA, Tukey post hoc test. ns= not significant, p>0.05). Data are all represented as mean values ± S.E.M.

**Supplemental Table.**
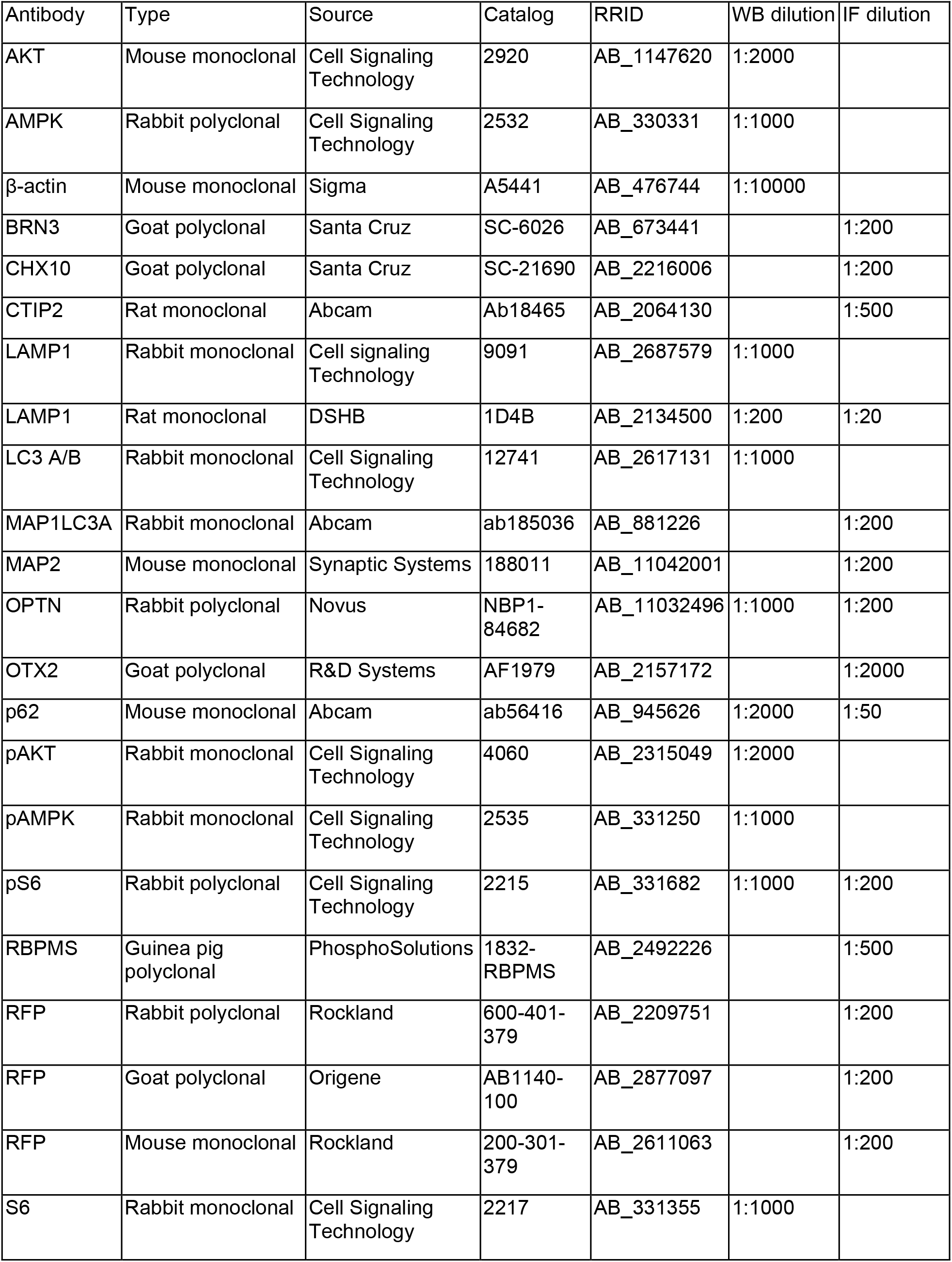
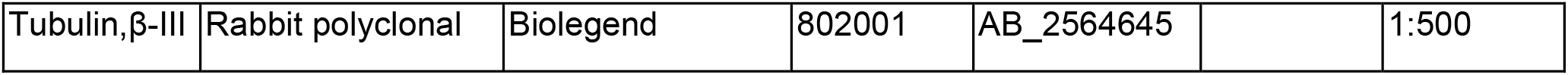
List of antibodies.

## References

1. Agostinone, J., Alarcon-Martinez, L., Gamlin, C., Yu, W.Q., Wong, R.O.L., and Di Polo, A. (2018). Insulin signalling promotes dendrite and synapse regeneration and restores circuit function after axonal injury. Brain 141, 1963–1980. 10.1093/brain/awy142.

2. Anborgh, P.H., Godin, C., Pampillo, M., Dhami, G.K., Dale, L.B., Cregan, S.P., Truant, R., and Ferguson, S.S. (2005). Inhibition of metabotropic glutamate receptor signaling by the huntingtin-binding protein optineurin. The Journal of biological chemistry 280, 34840–34848. 10.1074/jbc.M504508200.

3. Aung, T., Rezaie, T., Okada, K., Viswanathan, A.C., Child, A.H., Brice, G., Bhattacharya, S.S., Lehmann, O.J., Sarfarazi, M., and Hitchings, R.A. (2005). Clinical features and course of patients with glaucoma with the E50K mutation in the optineurin gene. Invest Ophthalmol Vis Sci 46, 2816–2822. 10.1167/iovs.04-1133.

4. Barmada, S.J., Serio, A., Arjun, A., Bilican, B., Daub, A., Ando, D.M., Tsvetkov, A., Pleiss, M., Li, X., Peisach, D., et al. (2014). Autophagy induction enhances TDP43 turnover and survival in neuronal ALS models. Nat Chem Biol 10, 677–685. 10.1038/nchembio.1563.

5. Belforte, N., Agostinone, J., Alarcon-Martinez, L., Villafranca-Baughman, D., Dotigny, F., Cueva Vargas, J.L., and Di Polo, A. (2021). AMPK hyperactivation promotes dendrite retraction, synaptic loss, and neuronal dysfunction in glaucoma. Mol Neurodegener 16, 43. 10.1186/s13024-021-00466-z.

6. Boya, P., Reggiori, F., and Codogno, P. (2013). Emerging regulation and functions of autophagy. Nat Cell Biol 15, 713–720. 10.1038/ncb2788.

7. Casadio, A., Martin, K.C., Giustetto, M., Zhu, H., Chen, M., Bartsch, D., Bailey, C.H., and Kandel, E.R. (1999). A transient, neuron-wide form of CREB-mediated long-term facilitation can be stabilized at specific synapses by local protein synthesis. Cell 99, 221–237. 10.1016/s0092-8674(00)81653-0.

8. Chalasani, M.L., Kumari, A., Radha, V., and Swarup, G. (2014). E50K-OPTN-induced retinal cell death involves the Rab GTPase-activating protein, TBC1D17 mediated block in autophagy. PLoS One 9, e95758. 10.1371/journal.pone.0095758.

9. Cuervo, A.M., Stefanis, L., Fredenburg, R., Lansbury, P.T., and Sulzer, D. (2004). Impaired degradation of mutant alpha-synuclein by chaperone-mediated autophagy. Science 305, 1292–1295. 10.1126/science.1101738.

10. De Marco, N., Buono, M., Troise, F., and Diez-Roux, G. (2006). Optineurin increases cell survival and translocates to the nucleus in a Rab8-dependent manner upon an apoptotic stimulus. The Journal of biological chemistry 281, 16147–16156. 10.1074/jbc.M601467200.

11. Duan, X., Qiao, M., Bei, F., Kim, I.J., He, Z., and Sanes, J.R. (2015). Subtype-specific regeneration of retinal ganglion cells following axotomy: effects of osteopontin and mTOR signaling. Neuron 85, 1244–1256. 10.1016/j.neuron.2015.02.017.

12. Evans, C.S., and Holzbaur, E.L. (2020). Degradation of engulfed mitochondria is rate-limiting in Optineurin-mediated mitophagy in neurons. Elife 9. 10.7554/eLife.50260.

13. Fligor, C.M., Huang, K.C., Lavekar, S.S., VanderWall, K.B., and Meyer, J.S. (2020). Differentiation of retinal organoids from human pluripotent stem cells. Methods Cell Biol 159, 279–302. 10.1016/bs.mcb.2020.02.005.

14. Ganley, I.G., Lam du, H., Wang, J., Ding, X., Chen, S., and Jiang, X. (2009). ULK1.ATG13.FIP200 complex mediates mTOR signaling and is essential for autophagy. J Biol Chem 284, 12297–12305. 10.1074/jbc.M900573200.

15. Guertin, D.A., and Sabatini, D.M. (2009). The pharmacology of mTOR inhibition. Sci Signal 2, pe24. 10.1126/scisignal.267pe24.

16. Hirt, J., Porter, K., Dixon, A., McKinnon, S., and Liton, P.B. (2018). Contribution of autophagy to ocular hypertension and neurodegeneration in the DBA/2J spontaneous glaucoma mouse model. Cell Death Discov 4, 14. 10.1038/s41420-018-0077-y.

17. Hosokawa, N., Hara, T., Kaizuka, T., Kishi, C., Takamura, A., Miura, Y., Iemura, S., Natsume, T., Takehana, K., Yamada, N., et al. (2009). Nutrient-dependent mTORC1 association with the ULK1-Atg13-FIP200 complex required for autophagy. Mol Biol Cell 20, 1981–1991. 10.1091/mbc.E08-12-1248.

18. Inoki, K., Kim, J., and Guan, K.L. (2012). AMPK and mTOR in cellular energy homeostasis and drug targets. Annu Rev Pharmacol Toxicol 52, 381–400. 10.1146/annurev-pharmtox-010611-134537.

19. Ito, Y.A., Belforte, N., Cueva Vargas, J.L., and Di Polo, A. (2016). A Magnetic Microbead Occlusion Model to Induce Ocular Hypertension-Dependent Glaucoma in Mice. J Vis Exp, e53731. 10.3791/53731.

20. Jefferies, H.B., Fumagalli, S., Dennis, P.B., Reinhard, C., Pearson, R.B., and Thomas, G. (1997). Rapamycin suppresses 5’TOP mRNA translation through inhibition of p70s6k. EMBO J 16, 3693–3704. 10.1093/emboj/16.12.3693.

21. Ji, M.M., Wang, L., Zhan, Q., Xue, W., Zhao, Y., Zhao, X., Xu, P.P., Shen, Y., Liu, H., Janin, A., et al. (2015). Induction of autophagy by valproic acid enhanced lymphoma cell chemosensitivity through HDAC-independent and IP3-mediated PRKAA activation. Autophagy 11, 2160–2171. 10.1080/15548627.2015.1082024.

22. Jung, S., Chung, Y., Lee, Y., Lee, Y., Cho, J.W., Shin, E.J., Kim, H.C., and Oh, Y.J. (2019). Buffering of cytosolic calcium plays a neuroprotective role by preserving the autophagy-lysosome pathway during MPP(+)-induced neuronal death. Cell Death Discov 5, 130. 10.1038/s41420-019-0210-6.

23. Kaizuka, T., Morishita, H., Hama, Y., Tsukamoto, S., Matsui, T., Toyota, Y., Kodama, A., Ishihara, T., Mizushima, T., and Mizushima, N. (2016). An Autophagic Flux Probe that Releases an Internal Control. Mol Cell 64, 835–849. 10.1016/j.molcel.2016.09.037.

24. Kim, D.H., Sarbassov, D.D., Ali, S.M., King, J.E., Latek, R.R., Erdjument-Bromage, H., Tempst, P., and Sabatini, D.M. (2002). mTOR interacts with raptor to form a nutrient-sensitive complex that signals to the cell growth machinery. Cell 110, 163–175. 10.1016/s0092-8674(02)00808-5.

25. Klionsky, D.J., Abdel-Aziz, A.K., Abdelfatah, S., Abdellatif, M., Abdoli, A., Abel, S., Abeliovich, H., Abildgaard, M.H., Abudu, Y.P., Acevedo-Arozena, A., et al. (2021). Guidelines for the use and interpretation of assays for monitoring autophagy (4th edition)(1). Autophagy 17, 1–382. 10.1080/15548627.2020.1797280.

26. Kulkarni, A., Dong, A., Kulkarni, V.V., Chen, J., Laxton, O., Anand, A., and Maday, S. (2020). Differential regulation of autophagy during metabolic stress in astrocytes and neurons. Autophagy 16, 1651–1667. 10.1080/15548627.2019.1703354.

27. Levine, B., and Kroemer, G. (2008). Autophagy in the pathogenesis of disease. Cell 132, 27–42. 10.1016/j.cell.2007.12.018.

28. Li, F., Xie, X., Wang, Y., Liu, J., Cheng, X., Guo, Y., Gong, Y., Hu, S., and Pan, L. (2016). Structural insights into the interaction and disease mechanism of neurodegenerative disease-associated optineurin and TBK1 proteins. Nat Commun 7, 12708. 10.1038/ncomms12708.

29. Linda, K., Lewerissa, E.I., Verboven, A.H.A., Gabriele, M., Frega, M., Klein Gunnewiek, T.M., Devilee, L., Ulferts, E., Hommersom, M., Oudakker, A., et al. (2022). Imbalanced autophagy causes synaptic deficits in a human model for neurodevelopmental disorders. Autophagy 18, 423–442. 10.1080/15548627.2021.1936777.

30. Lipton, J.O., and Sahin, M. (2014). The neurology of mTOR. Neuron 84, 275–291. 10.1016/j.neuron.2014.09.034.

31. Liu, X., Wang, Q., Shao, Z., Zhang, S., Hou, M., Jiang, M., Du, M., Li, J., and Yuan, H. (2021). Proteomic analysis of aged and OPTN E50K retina in the development of normal tension glaucoma. Hum Mol Genet 30, 1030–1044. 10.1093/hmg/ddab099.

32. Livak, K.J., and Schmittgen, T.D. (2001). Analysis of relative gene expression data using real-time quantitative PCR and the 2(-Delta Delta C(T)) Method. Methods 25, 402–408. 10.1006/meth.2001.1262.

33. Maday, S., and Holzbaur, E.L. (2016). Compartment-Specific Regulation of Autophagy in Primary Neurons. J Neurosci 36, 5933–5945. 10.1523/JNEUROSCI.4401-15.2016.

34. Madrakhimov, S.B., Yang, J.Y., Kim, J.H., Han, J.W., and Park, T.K. (2021). mTOR-dependent dysregulation of autophagy contributes to the retinal ganglion cell loss in streptozotocin-induced diabetic retinopathy. Cell Commun Signal 19, 29. 10.1186/s12964-020-00698-4.

35. Markovinovic, A., Ljutic, T., Beland, L.C., and Munitic, I. (2018). Optineurin Insufficiency Disbalances Proinflammatory and Anti-inflammatory Factors by Reducing Microglial IFN-beta Responses. Neuroscience 388, 139–151. 10.1016/j.neuroscience.2018.07.007.

36. Maruyama, H., Morino, H., Ito, H., Izumi, Y., Kato, H., Watanabe, Y., Kinoshita, Y., Kamada, M., Nodera, H., Suzuki, H., et al. (2010). Mutations of optineurin in amyotrophic lateral sclerosis. Nature 465, 223–226. 10.1038/nature08971.

37. Menzies, F.M., Fleming, A., and Rubinsztein, D.C. (2015). Compromised autophagy and neurodegenerative diseases. Nat Rev Neurosci 16, 345–357. 10.1038/nrn3961.

38. Meyer, J.S., Howden, S.E., Wallace, K.A., Verhoeven, A.D., Wright, L.S., Capowski, E.E., Pinilla, I., Martin, J.M., Tian, S., Stewart, R., et al. (2011). Optic vesicle-like structures derived from human pluripotent stem cells facilitate a customized approach to retinal disease treatment. Stem Cells 29, 1206–1218. 10.1002/stem.674.

39. Minegishi, Y., Iejima, D., Kobayashi, H., Chi, Z.L., Kawase, K., Yamamoto, T., Seki, T., Yuasa, S., Fukuda, K., and Iwata, T. (2013). Enhanced optineurin E50K-TBK1 interaction evokes protein insolubility and initiates familial primary open-angle glaucoma. Hum Mol Genet 22, 3559–3567. 10.1093/hmg/ddt210.

40. Mori, F., Tanji, K., Toyoshima, Y., Yoshida, M., Kakita, A., Takahashi, H., and Wakabayashi, K. (2012). Optineurin immunoreactivity in neuronal nuclear inclusions of polyglutamine diseases (Huntington’s, DRPLA, SCA2, SCA3) and intranuclear inclusion body disease. Acta Neuropathol 123, 747–749. 10.1007/s00401-012-0956-x.

41. Munitic, I., Giardino Torchia, M.L., Meena, N.P., Zhu, G., Li, C.C., and Ashwell, J.D. (2013). Optineurin insufficiency impairs IRF3 but not NF-kappaB activation in immune cells. J Immunol 191, 6231–6240. 10.4049/jimmunol.1301696.

42. Nixon, R.A. (2013). The role of autophagy in neurodegenerative disease. Nat Med 19, 983–997. 10.1038/nm.3232.

43. Nixon, R.A., and Yang, D.S. (2012). Autophagy and neuronal cell death in neurological disorders. Cold Spring Harb Perspect Biol 4. 10.1101/cshperspect.a008839.

44. Ohlemacher, S.K., Iglesias, C.L., Sridhar, A., Gamm, D.M., and Meyer, J.S. (2015). Generation of highly enriched populations of optic vesicle-like retinal cells from human pluripotent stem cells. Curr Protoc Stem Cell Biol 32, 1H 8 1–1H 8 20. 10.1002/9780470151808.sc01h08s32.

45. Osawa, T., Mizuno, Y., Fujita, Y., Takatama, M., Nakazato, Y., and Okamoto, K. (2011). Optineurin in neurodegenerative diseases. Neuropathology 31, 569–574. 10.1111/j.1440-1789.2011.01199.x.

46. Palmieri, M., Pal, R., Nelvagal, H.R., Lotfi, P., Stinnett, G.R., Seymour, M.L., Chaudhury, A., Bajaj, L., Bondar, V.V., Bremner, L., et al. (2017). mTORC1-independent TFEB activation via Akt inhibition promotes cellular clearance in neurodegenerative storage diseases. Nat Commun 8, 14338. 10.1038/ncomms14338.

47. Park, B.C., Shen, X., Samaraweera, M., and Yue, B.Y. (2006). Studies of optineurin, a glaucoma gene: Golgi fragmentation and cell death from overexpression of wild-type and mutant optineurin in two ocular cell types. Am J Pathol 169, 1976–1989. 10.2353/ajpath.2006.060400.

48. Park, K.K., Liu, K., Hu, Y., Smith, P.D., Wang, C., Cai, B., Xu, B., Connolly, L., Kramvis, I., Sahin, M., and He, Z. (2008). Promoting axon regeneration in the adult CNS by modulation of the PTEN/mTOR pathway. Science 322, 963–966. 10.1126/science.1161566.

49. Pernet, V., Bourgeois, P., and Di Polo, A. (2007). A role for polyamines in retinal ganglion cell excitotoxic death. J Neurochem 103, 1481–1490. 10.1111/j.1471-4159.2007.04843.x.

50. Porter, K., Hirt, J., Stamer, W.D., and Liton, P.B. (2015). Autophagic dysregulation in glaucomatous trabecular meshwork cells. Biochim Biophys Acta 1852, 379–385. 10.1016/j.bbadis.2014.11.021.

51. Porter, K.M., Jeyabalan, N., and Liton, P.B. (2014). MTOR-independent induction of autophagy in trabecular meshwork cells subjected to biaxial stretch. Biochim Biophys Acta 1843, 1054–1062. 10.1016/j.bbamcr.2014.02.010.

52. Proietti Onori, M., Koene, L.M.C., Schäfer, C.B., Nellist, M., de Brito van Velze, M., Gao, Z., Elgersma, Y., and van Woerden, G.M. (2021). RHEB/mTOR hyperactivity causes cortical malformations and epileptic seizures through increased axonal connectivity. PLoS Biol 19, e3001279. 10.1371/journal.pbio.3001279.

53. Qassim, A., and Siggs, O.M. (2020). Predicting the genetic risk of glaucoma. The Biochemist 42, 26–30.

54. Qiu, Y., Wang, J., Li, H., Yang, B., Wang, J., He, Q., and Weng, Q. (2021). Emerging views of OPTN (optineurin) function in the autophagic process associated with disease. Autophagy, 1–13. 10.1080/15548627.2021.1908722.

55. Querfurth, H., and Lee, H.K. (2021). Mammalian/mechanistic target of rapamycin (mTOR) complexes in neurodegeneration. Mol Neurodegener 16, 44. 10.1186/s13024-021-00428-5.

56. Rezaie, T., Child, A., Hitchings, R., Brice, G., Miller, L., Coca-Prados, M., Heon, E., Krupin, T., Ritch, R., Kreutzer, D., et al. (2002). Adult-onset primary open-angle glaucoma caused by mutations in optineurin. Science 295, 1077–1079. 10.1126/science.1066901.

57. Ritch, R., Darbro, B., Menon, G., Khanna, C.L., Solivan-Timpe, F., Roos, B.R., Sarfarzi, M., Kawase, K., Yamamoto, T., Robin, A.L., et al. (2014). TBK1 gene duplication and normal-tension glaucoma. JAMA Ophthalmol 132, 544–548. 10.1001/jamaophthalmol.2014.104.

58. Rusmini, P., Cortese, K., Crippa, V., Cristofani, R., Cicardi, M.E., Ferrari, V., Vezzoli, G., Tedesco, B., Meroni, M., Messi, E., et al. (2019). Trehalose induces autophagy via lysosomal-mediated TFEB activation in models of motoneuron degeneration. Autophagy 15, 631–651. 10.1080/15548627.2018.1535292.

59. Schwab, C., Yu, S., McGeer, E.G., and McGeer, P.L. (2012). Optineurin in Huntington’s disease intranuclear inclusions. Neurosci Lett 506, 149–154. 10.1016/j.neulet.2011.10.070.

60. Shen, W.C., Li, H.Y., Chen, G.C., Chern, Y., and Tu, P.H. (2015). Mutations in the ubiquitin-binding domain of OPTN/optineurin interfere with autophagy-mediated degradation of misfolded proteins by a dominant-negative mechanism. Autophagy 11, 685–700. 10.4161/auto.36098.

61. Shim, M.S., Takihara, Y., Kim, K.Y., Iwata, T., Yue, B.Y., Inatani, M., Weinreb, R.N., Perkins, G.A., and Ju, W.K. (2016). Mitochondrial pathogenic mechanism and degradation in optineurin E50K mutation-mediated retinal ganglion cell degeneration. Sci Rep 6, 33830. 10.1038/srep33830.

62. Shoemaker, C.J., Huang, T.Q., Weir, N.R., Polyakov, N.J., Schultz, S.W., and Denic, V. (2019). CRISPR screening using an expanded toolkit of autophagy reporters identifies TMEM41B as a novel autophagy factor. PLoS Biol 17, e2007044. 10.1371/journal.pbio.2007044.

63. Singh, K., Matsuyama, S., Drazba, J.A., and Almasan, A. (2012). Autophagy-dependent senescence in response to DNA damage and chronic apoptotic stress. Autophagy 8, 236–251. 10.4161/auto.8.2.18600.

64. Sirohi, K., and Swarup, G. (2016). Defects in autophagy caused by glaucoma-associated mutations in optineurin. Experimental eye research 144, 54–63. 10.1016/j.exer.2015.08.020.

65. Slowicka, K., Vereecke, L., Mc Guire, C., Sze, M., Maelfait, J., Kolpe, A., Saelens, X., Beyaert, R., and van Loo, G. (2016). Optineurin deficiency in mice is associated with increased sensitivity to Salmonella but does not affect proinflammatory NF-kappaB signaling. Eur J Immunol 46, 971–980. 10.1002/eji.201545863.

66. Sluch, V.M., Chamling, X., Liu, M.M., Berlinicke, C.A., Cheng, J., Mitchell, K.L., Welsbie, D.S., and Zack, D.J. (2017). Enhanced Stem Cell Differentiation and Immunopurification of Genome Engineered Human Retinal Ganglion Cells. Stem Cells Transl Med 6, 1972–1986. 10.1002/sctm.17-0059.

67. Swarup, G., and Sayyad, Z. (2018). Altered Functions and Interactions of Glaucoma-Associated Mutants of Optineurin. Front Immunol 9, 1287. 10.3389/fimmu.2018.01287.

68. Tang, G., Gudsnuk, K., Kuo, S.H., Cotrina, M.L., Rosoklija, G., Sosunov, A., Sonders, M.S., Kanter, E., Castagna, C., Yamamoto, A., et al. (2014). Loss of mTOR-dependent macroautophagy causes autistic-like synaptic pruning deficits. Neuron 83, 1131–1143. 10.1016/j.neuron.2014.07.040.

69. Tavazoie, S.F., Alvarez, V.A., Ridenour, D.A., Kwiatkowski, D.J., and Sabatini, B.L. (2005). Regulation of neuronal morphology and function by the tumor suppressors Tsc1 and Tsc2. Nat Neurosci 8, 1727–1734. 10.1038/nn1566.

70. Teotia, P., Van Hook, M.J., Fischer, D., and Ahmad, I. (2019). Human retinal ganglion cell axon regeneration by recapitulating developmental mechanisms: effects of recruitment of the mTOR pathway. Development 146. 10.1242/dev.178012.

71. Tseng, H.C., Riday, T.T., McKee, C., Braine, C.E., Bomze, H., Barak, I., Marean-Reardon, C., John, S.W., Philpot, B.D., and Ehlers, M.D. (2015). Visual impairment in an optineurin mouse model of primary open-angle glaucoma. Neurobiol Aging 36, 2201–2212. 10.1016/j.neurobiolaging.2015.02.012.

72. VanderWall, K.B., Huang, K.C., Pan, Y., Lavekar, S.S., Fligor, C.M., Allsop, A.R., Lentsch, K.A., Dang, P., Zhang, C., Tseng, H.C., et al. (2020). Retinal Ganglion Cells With a Glaucoma OPTN(E50K) Mutation Exhibit Neurodegenerative Phenotypes when Derived from Three-Dimensional Retinal Organoids. Stem Cell Reports 15, 52–66. 10.1016/j.stemcr.2020.05.009.

73. Wareham, L.K., Liddelow, S.A., Temple, S., Benowitz, L.I., Di Polo, A., Wellington, C., Goldberg, J.L., He, Z., Duan, X., Bu, G., et al. (2022). Solving neurodegeneration: common mechanisms and strategies for new treatments. Mol Neurodegener 17, 23. 10.1186/s13024-022-00524-0.

74. White, K.E., Davies, V.J., Hogan, V.E., Piechota, M.J., Nichols, P.P., Turnbull, D.M., and Votruba, M. (2009). OPA1 deficiency associated with increased autophagy in retinal ganglion cells in a murine model of dominant optic atrophy. Invest Ophthalmol Vis Sci 50, 2567–2571. 10.1167/iovs.08-2913.

75. Wong, Y.C., and Holzbaur, E.L. (2014). Optineurin is an autophagy receptor for damaged mitochondria in parkin-mediated mitophagy that is disrupted by an ALS-linked mutation. Proc Natl Acad Sci U S A 111, E4439–4448. 10.1073/pnas.1405752111.

76. Wong, Y.C., and Holzbaur, E.L. (2015). Autophagosome dynamics in neurodegeneration at a glance. J Cell Sci 128, 1259–1267. 10.1242/jcs.161216.

77. Yang, Y., Wang, Q., Song, D., Zen, R., Zhang, L., Wang, Y., Yang, H., Zhang, D., Jia, J., Zhang, J., and Wang, J. (2020). Lysosomal dysfunction and autophagy blockade contribute to autophagy-related cancer suppressing peptide-induced cytotoxic death of cervical cancer cells through the AMPK/mTOR pathway. J Exp Clin Cancer Res 39, 197. 10.1186/s13046-020-01701-z.

78. Ying, H., and Yue, B.Y. (2012). Cellular and molecular biology of optineurin. Int Rev Cell Mol Biol 294, 223–258. 10.1016/B978-0-12-394305-7.00005-7.

79. Yu, W.H., Cuervo, A.M., Kumar, A., Peterhoff, C.M., Schmidt, S.D., Lee, J.H., Mohan, P.S., Mercken, M., Farmery, M.R., Tjernberg, L.O., et al. (2005). Macroautophagy--a novel Beta-amyloid peptide-generating pathway activated in Alzheimer’s disease. J Cell Biol 171, 87–98. 10.1083/jcb.200505082.

